# The depletion of TRAIP results in the retention of PCNA on chromatin during mitosis, leads to inhibiting DNA replication initiation

**DOI:** 10.1101/2023.05.08.539800

**Authors:** Yasushi Shiomi, Akiyo Hayashi, Yuichiro Saito, Masato T. Kanemaki, Hideo Nishitani

**Author notes:** Molecular Biology Program, Memorial Sloan Kettering Cancer Center, New York, NY 10065, USA.

## Abstract

Loading PCNA onto chromatin is a pivotal step in DNA replication, cell cycle progression, and genome integrity. Conversely, unloading PCNA from chromatin is equally crucial for maintaining genome stability. Cells deficient in the PCNA unloader ATAD5-RFC exhibit elevated levels of chromatin-bound PCNA during S phase, but still show dissociation of PCNA from chromatin in mitosis. In this study, we found that depletion of TRAIP, an E3 ubiquitin ligase, results in the retention of PCNA on chromatin during mitosis. Although TRAIP-depleted cells with chromatin-bound PCNA during mitosis progressed into the subsequent G1 phase, they displayed reduced levels of Cdt1, a key replication licensing factor, and impaired S phase entry. In addition, TRAIP-depleted cells exhibited delayed S phase progression. These results suggest that TRAIP functions independently of ATAD5-RFC in removing PCNA from chromatin. Furthermore, TRAIP appears to be essential for precise pre-replication complexes (pre-RCs) formation necessary for faithful initiation of DNA replication and S phase progression.

## INTRODUCTION

DNA replication initiation involves origin licensing during late mitosis and G1 phase. This process forms pre-replication complexes (pre-RCs) by loading MCM2-7 onto ORC-bound origins, facilitated by Cdc6 and Cdt1. Cdt1, a key licensing regulator, is rapidly degraded by CRL1-Skp2 and CRL4-Cdt2 E3 ubiquitin ligases after replication onset to prevent re-licensing. CDK and DDK activate pre-RCs, leading to CMG (Cdc45-MCM2-7-GINS) helicase complex formation (Nishitani and Lygerou 2004; Masai et al. 2010; Havens and Walter 2011; Costa and Diffley 2022). Following origin unwinding and primer synthesis, DNA polymerases bind to PCNA for processive replication. Replication factor C (RFC) complex loads PCNA onto primer/template junctions. PCNA is essential for DNA synthesis and coordinates various replication-linked processes (Waga and Stillman 1998; Kelch et al. 2012; Burgers and Kunkel 2017).

In addition to an authentic RFC complex, RFC1-RFC, eukaryotic cells possess multiple RFC complexes: Ctf18-RFC, and ATAD5-RFC (Elg1-RFC in yeast) (Shiomi and Nishitani 2017; Ohashi and Tsurimoto 2017). RNAi knockdown studies revealed that ATAD5 depletion results in elevated chromatin-bound PCNA levels, delayed S phase progression, abnormal chromatin protein dynamics, and aberrant chromosome structures. These results demonstrated that unlike other RFC complexes, ATAD5-RFC plays a crucial role in maintaining genome stability by regulating PCNA unloading from chromatin (Lee et al. 2013; Kubota et al. 2013; Shiomi and Nishitani 2013).

When DNA synthesis during replication is terminated, CMG disassembly is triggered by MCM7 ubiquitination (Maric et al. 2014; Moreno et al. 2014). During S phase, SCF-Dia2 in *S. cerevisiae* (Maric et al. 2017) and CUL2-LRR1 in *C. elegans* (Merlet et al. 2010; Burger et al. 2013; Sonneville et al. 2017) and *X. laevis* (Dewar et al. 2017; Sonneville et al. 2017) ubiquitinate MCM7. The AAA+ ATPase p97/VCP/Cdc48 then unfolds ubiquitinated MCM7, irreversibly dissociating the CMG complex from chromatin (Maric et al. 2017; Mukherjee and Labib 2019; Deegan et al. 2020). In LRR-1-depleted cells, CMG persists on chromatin after replication but dissociates upon mitotic entry. Mitotic CMG disassembly requires the a metazoan-specific RING ubiquitin ligase known as TRUL-1 in *C. elegans* and TRAIP in vertebrates (Sonneville et al. 2017; Deng et al. 2019; Priego Moreno et al. 2019; Sonneville et al. 2019). TRAIP is essential for CMG disassembly during mitosis, at stalled replisomes prior to mitotic DNA synthesis (MiDAS), and at sites of interstrand crosslinks (ICLs) or replication-transcription conflicts in S phase. TRAIP also facilitates DNA damage signaling and repair by interacting with various repair factors involved in the DNA damage response (Chapard et al. 2014; Wallace et al. 2014; Harley et al. 2016; Hoffmann et al. 2016; Soo Lee et al. 2016; Han et al. 2019; Sonneville et al. 2019; Li et al. 2020; Wu, Pellman and Walter 2020; Scaramuzza et al., 2023).

Here, we propose a novel function of TRAIP in removing PCNA from chromatin during normal cell cycle progression. TRAIP-depleted cells exhibited retention of PCNA on chromatin throughout mitosis. This persistent chromatin-bound PCNA induced Cdt1 degradation during the subsequent G1 phase, impairing pre-RCs formation. Consequently, S phase entry was compromised, resulting in delayed S phase progression.

## RESULTS and DISCUSSION

### TRAIP mediated dissociation of PCNA from chromatin during normal cell cycle progression

Consistent with our previous findings (Shiomi and Nishitani, 2013), HCT116 cells expressing Tet-OsTIR1 and mAID-tagged ATAD5 exhibited elevated levels of chromatin-bound PCNA in asynchronous cells following doxycycline (dox) and auxin (IAA) treatment compared to control cells (Figure 1a). However, PCNA dissociated from chromatin even in ATAD5-depleted cells when arrested in mitosis, suggesting that cells possess additional mechanism for the removal of PCNA from chromatin.

**Figure 1.**
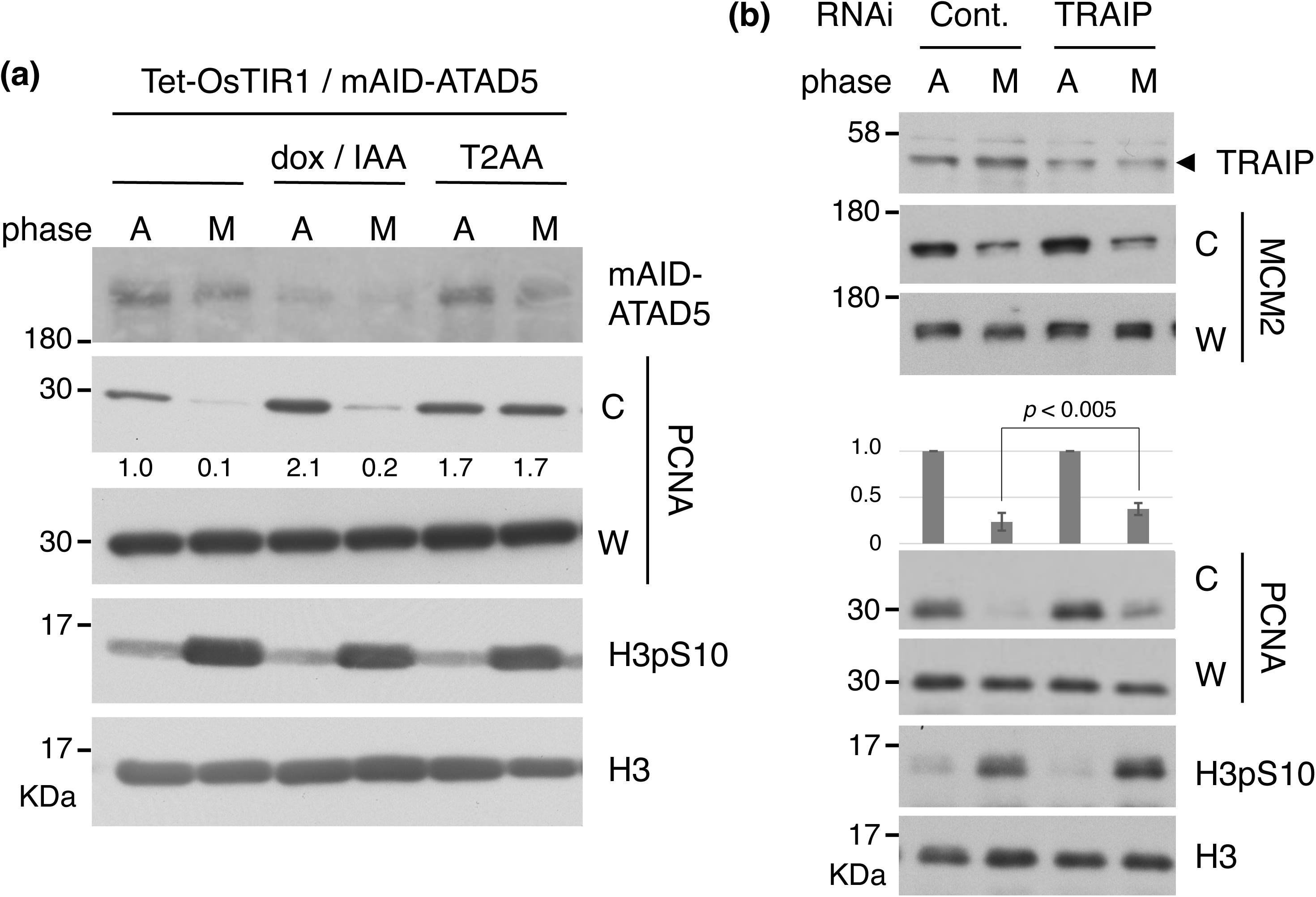
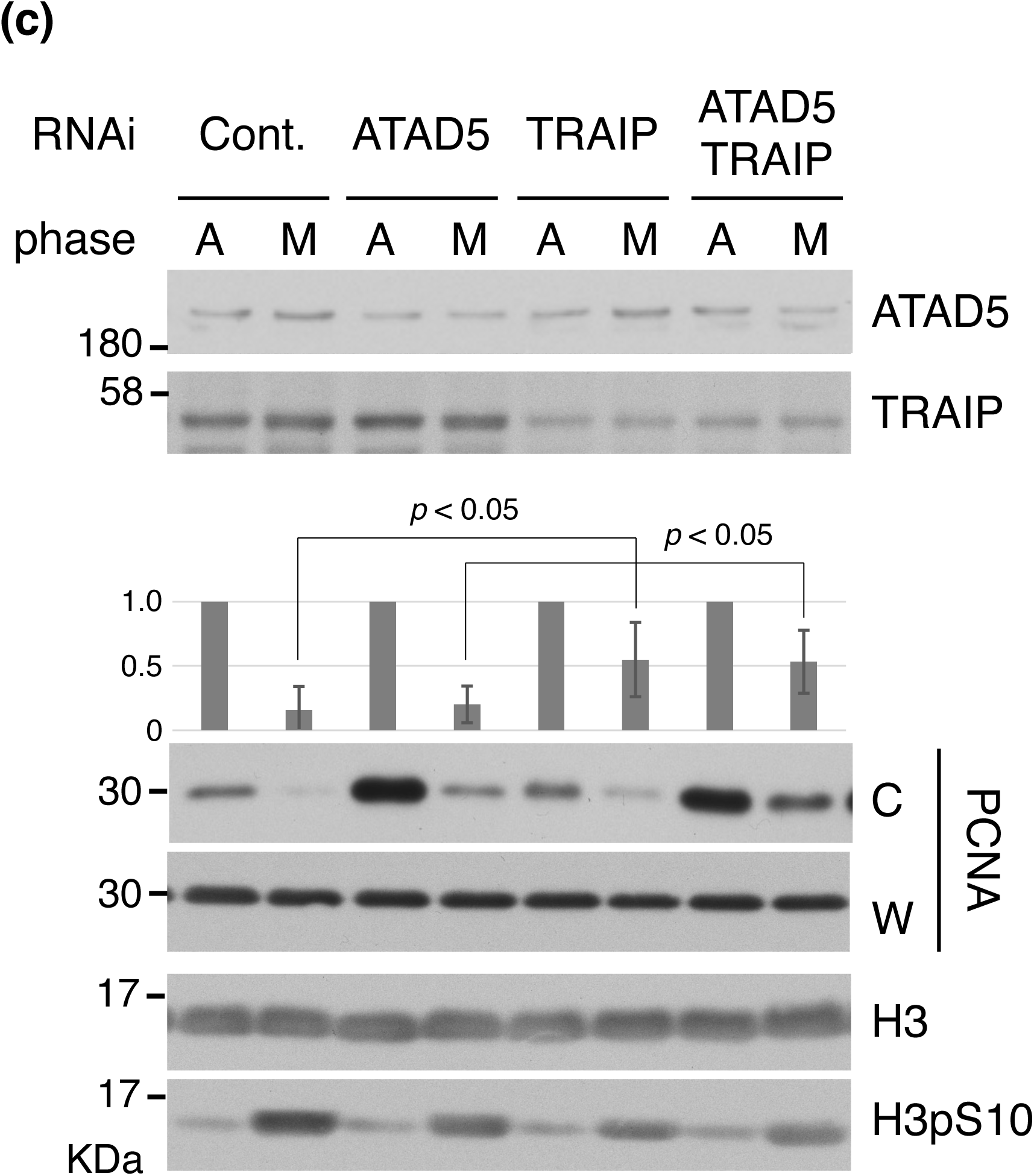
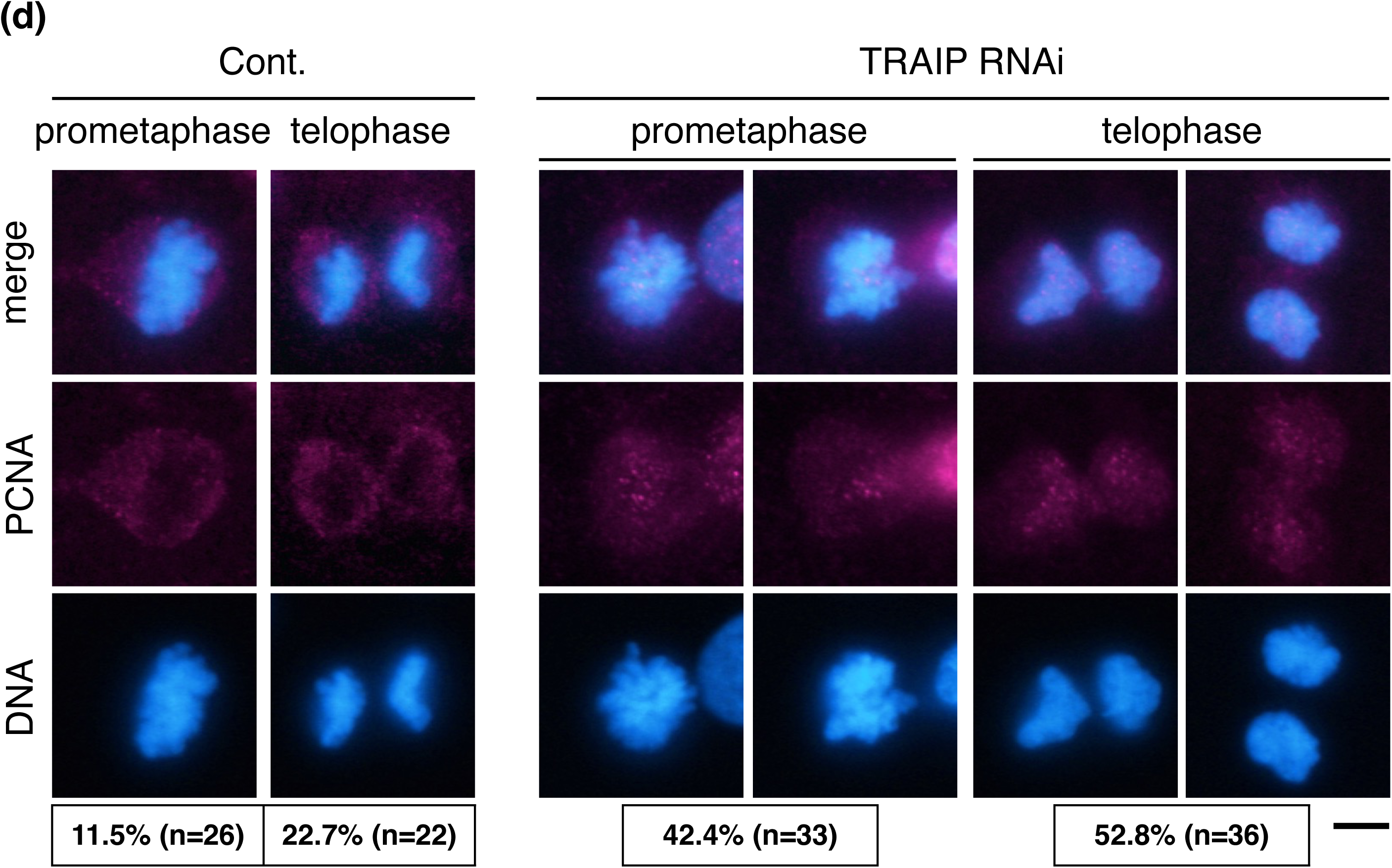
PCNA was removed from chromatin by TRAIP. (a) Asynchronous (phase; A) and mitosis-arrested (phase; M) HCT116 cells expressing mAID-ATAD5 were lysed and separated into whole cell extract fractions (W) and chromatin fractions (C), and the fractions were immunoblotted using the indicated antibodies. To deplete mAID-ATAD5, cells were cultured in medium with doxycycline and IAA for 24 hours, followed by 24 hours treatment with nocodazole to arrest the cells in mitosis, also in the presence of doxycycline and IAA. T2AA was added for 24 hours to inhibit the interaction between PCNA and PIP-containing proteins, followed by treatment with nocodazole for 24 hours to arrest the cells in mitosis, also in the presence of T2AA. The levels of chromatin-bound PCNA, normalized to the total PCNA levels in the whole cell extract (W), are indicated below the immunoblot bands, relative to those in asynchronous control cells set as 1.0. (b) Asynchronous (A) and mitosis-arrested (M) HCT116 cells with and without TRAIP depletion by RNAi were lysed and separated into W and C fractions, and the fractions were immunoblotted with the indicated antibodies. (c) Asynchronous (A) and mitosis-arrested (M) U2OS cells with and without ATAD5 and TRAIP depletion by RNAi were lysed and separated into W and C fractions, and the fractions were immunoblotted with the indicated antibodies. (d) Endogenous PCNA immunofluorescence in asynchronously growing U2OS cells, with or without TRAIP depletion by RNAi. The percentages of cells displaying at least one PCNA signal (cyan) on mitotic chromatin are indicated at the bottom of each image. The bar indicates 10 μm. The graphs (b and c) represent the averages of four independent experiments with standard deviations. The levels of chromatin-bound PCNA, normalized to total PCNA levels in the whole-cell extract (W), are displayed as vertical bars with error bars, relative to those in asynchronous control cells, which are set to 1.0. Unpaired two-sided Student’s t-tests were performed, and the corresponding p-values are indicated in the figures.

Based on previous reports (Hoffmann et al. 2016), it is conceivable that TRAIP associates with and ubiquitinates PCNA, potentially leading to its removal from chromatin, given the presence of a PCNA-interacting peptide sequence (PIP box) at the C-terminal end of TRAIP. Indeed, PCNA is retained on chromatin in mitotic-arrested cells treated with T2AA, an inhibitor of interactions between PCNA and PIP box-containing proteins (Punchihewa et al. 2012). We observed PCNA retention on chromatin during mitosis in TRAIP-depleted HCT116 cells, while chromatin-bound PCNA decreased in control cells (Figure 1b). Additionally, when comparing between control and ATAD5-depleted U2OS cells, as well as between cells depleted of TRAIP alone and cells depleted of both ATAD5 and TRAIP simultaneously, a similar reduction rates in chromatin-bound PCNA during mitosis was observed. This suggests that ATAD5-RFC and TRAIP function independently at different cell cycle stages (Figure 1c). Since nocodazole inhibits tubulin polymerization, cells are arrested at early mitosis. To determine whether PCNA was retained on chromatin throughout mitosis in TRAIP-depleted cells, we performed immunofluorescent detection of PCNA in asynchronously growing U2OS cells (Figure 1d). In control cells, PCNA signals were absent from chromatin during early (prometaphase) and late (telophase) mitosis. However, in TRAIP-depleted cells, PCNA signals frequently co-localized with chromatin throughout mitosis.

A previous study suggested PCNA as a candidate substrate for TRAIP (Hoffmann et al., 2016), and TRAIP-mediated regulation of chromatin-bound PCNA levels has been reported (Feng et al., 2016). However, that study was limited to TRAIP-mediated release of chromatin-bound PCNA during DNA damage-induced replication stress. Our results demonstrate that TRAIP is indispensable for PCNA removal from chromatin during normal cell cycle progression, providing new insights into the control of PCNA levels on chromatin.

### TRAIP was required for accurate cell cycle progression and correct cell growth

We previously reported that ATAD5-depleted cells exhibit a significant delay in S phase progression due to elevated levels of chromatin-bound PCNA (Shiomi and Nishitani 2013) (Figure 2a). When we analyzed the cell cycle profile of asynchronously growing TRAIP-depleted U2OS cells, we observed alterations across all phases of the cell cycle compared to control cells. To further elucidate these effects, we performed live-cell imaging of U2OS cells expressing EGFP-PCNA and depleted of TRAIP via RNAi (Figure S1, S2, S3 and S4). Comparison of the durations of each cell cycle phases between TRAIP-depleted and control cells revealed that TRAIP-depleted cells exhibited a prolonged G2 phase (Figure 2b). In contrast, no significant extension was observed in mitosis; however, a slight reduction in mitotic duration was noted on average. These results are consistent with the findings reported by Scaramuzza et al. and Chapard et al., respectively (Chapard et al., 2014; Scaramuzza et al., 2023). Furthermore, our analysis revealed that TRAIP-depleted cells exhibited a significant prolongation of S phase. In contrast, no substantial alteration was observed in the duration of G1 phase.

**Figure 2.**
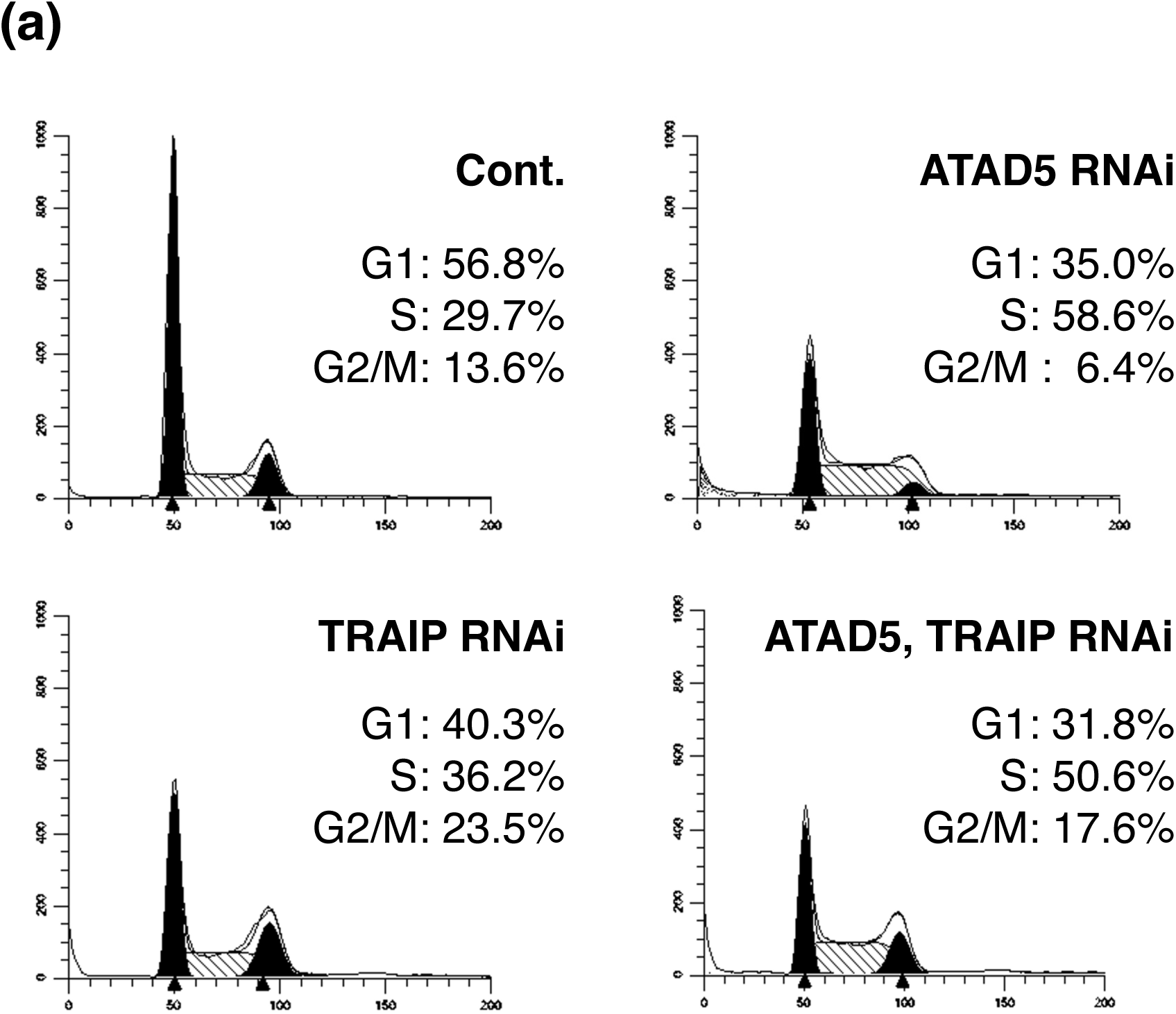
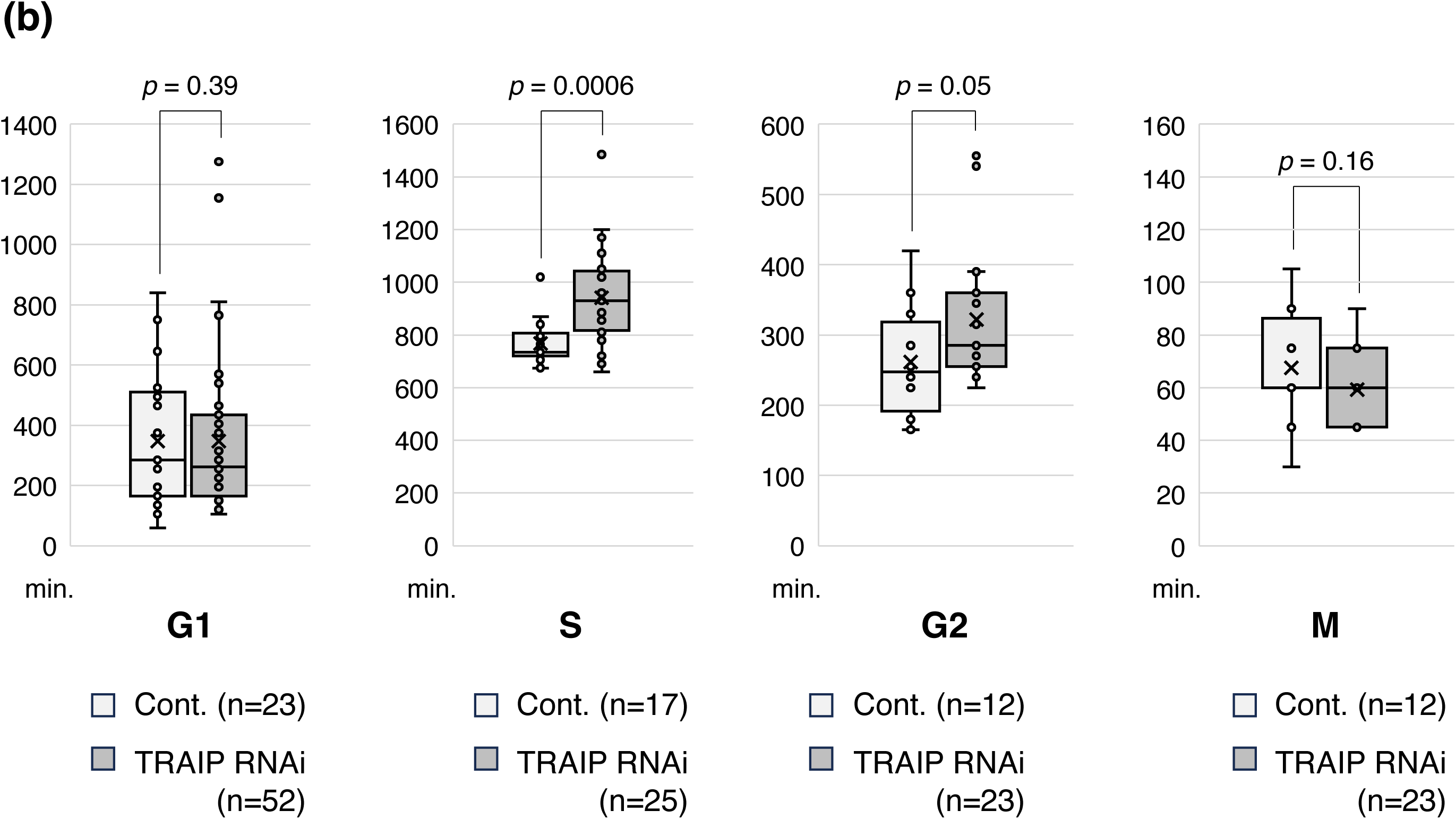
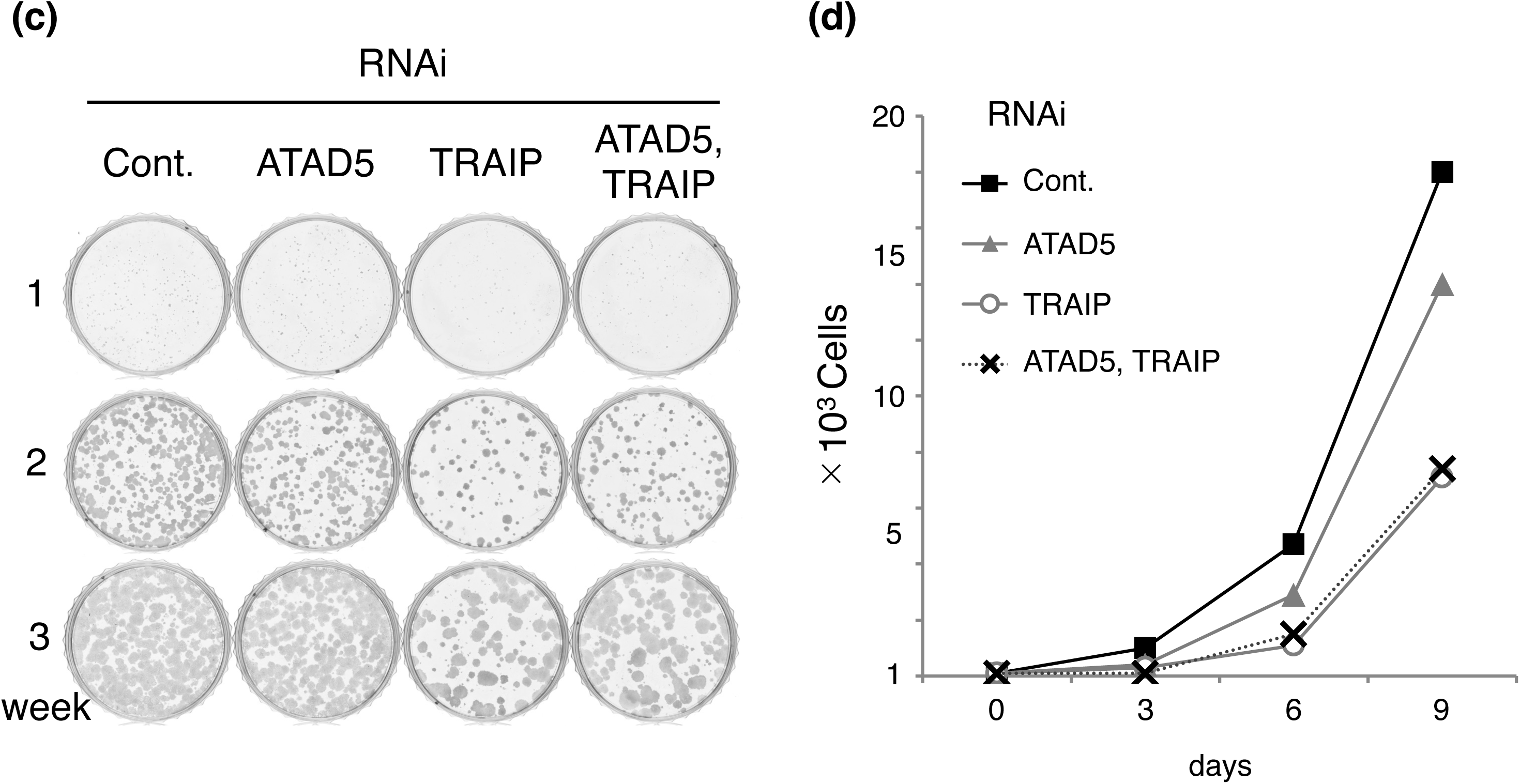
TRAIP is required for accurate cell cycle progression and appropriate cell growth. (a) Cell cycle profiles of asynchronously growing cells after depletion of the indicated proteins by RNAi. The ratios of cell populations at each cell cycle phase are indicated. (b) The durations of each cell cycle phases were determined from time-lapse images of asynchronously growing U2OS cells expressing GFP-PCNA, captured at 15-minute intervals, with and without TRAIP depletion by RNAi (Figure S1, S2, S3 and S4). The graph shows box plots of the indicated number of independent cells results. Unpaired two-sided Student’s t-tests were performed, and the corresponding p-values are indicated in the figure. The cross marks represent the mean values. The mean values for the time required for each phase progression (minutes) are as follows: TRAIP RNAi (G1 phase; average=351.2, median=262.5, S phase; average=941.4, median=930.0, G2 phase; average=310.7, median=285.0, mitosis; average=60.0, median=60.0), control RNAi (G1 phase; average=358.6, median=285.0, S phase; average=766.9, median=735.0, G2 phase; average=261.3, median=247.5, mitosis; average=67.5, median=60.0)). (c) Colony formation assays of U2OS cells with depletion of the indicated proteins by RNAi. The cells were cultured for indicated weeks. Colonies were stained with Coomassie Brilliant Blue (CBB). (d) Cell proliferation assays of U2OS cells with depletion of the indicated proteins by RNAi. The cells were cultured for indicated days. Cell numbers were determined using a hemocytometer.

The cell cycle progression defect in TRAIP-depleted cells was not due to activation of DNA replication, damage, or spindle assembly checkpoint, because activation of Chk1 and γH2AX and chromatin-bound Mad2 were not detected in TRAIP-depleted cells (Figure S5). A colony formation assay and measurement of the proliferation rate also showed that TRAIP is critical for correct cell growth (Figure 2c and 2d). Colonies formation and the cell growth of TRAIP-depleted cells was almost half that of control cells. ATAD5 and TRAIP double depleted cells exhibited almost the same growth and survival rate as TRAIP only depleted cells, indicating that TRAIP is more crucial for cell cycle progression than ATAD5-RFC. These results suggest that TRAIP is essential for precise cell proliferation, particularly during S phase progression.

### Clearance of PCNA from chromatin by TRAIP was required for DNA replication initiation

Next, we expressed AID2 technology based TRAIP tagged with mAID at its C-terminus in cells to analyze further TRAIP functions (Figure S6). The cell cycle profiles of TRAIP-mAID expressed cells up to 24 hours after auxin addition revealed a gradual increase in the 4C DNA content cell population, which is consistent with the result of the live-cell imaging experiment (Figure 3a and 2b). After 24 hours, cells that exhibited delayed mitotic exit as a result of prolonged G2 phase progression ultimately entered the G1 phase. However, their subsequent progression into the S phase appeared to be impaired. To validate the role of TRAIP in removal of PCNA from chromatin and cell cycle progression, we introduced untagged TRAIP cDNA constructs (wild-type and PIP box deletion mutant; Figure S7) into cells expressing TRAIP-mAID. As expected, cells expressing TRAIP (WT) exhibited reduced chromatin-bound PCNA upon mitotic synchronization following auxin addition. In contrast, cells expressing TRAIP (ΔPIP) maintained chromatin-bound PCNA under identical conditions (Figure S8). Furthermore, TRAIP (WT) attenuated alterations in cell cycle phase distributions relative to auxin-untreated control cells. Conversely, cells expressing TRAIP (ΔPIP) demonstrated cell cycle alterations similar to those observed upon TRAIP depletion (Figure S9). These findings demonstrate that TRAIP and its PIP box are essential for facilitating stable cell cycle progression and subsequent S phase entry through its functional activity, which involves the removal of PCNA from chromatin.

**Figure 3.**
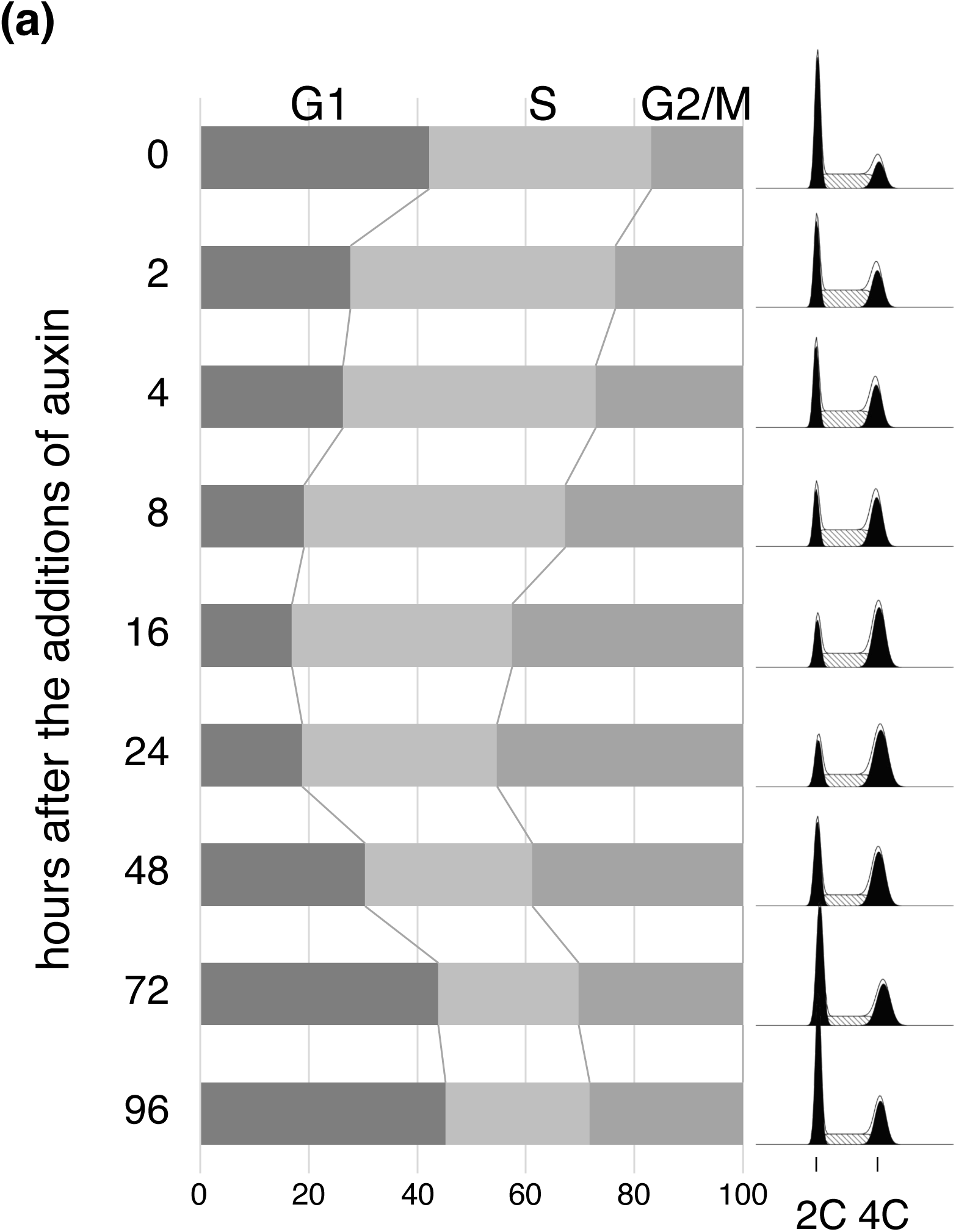
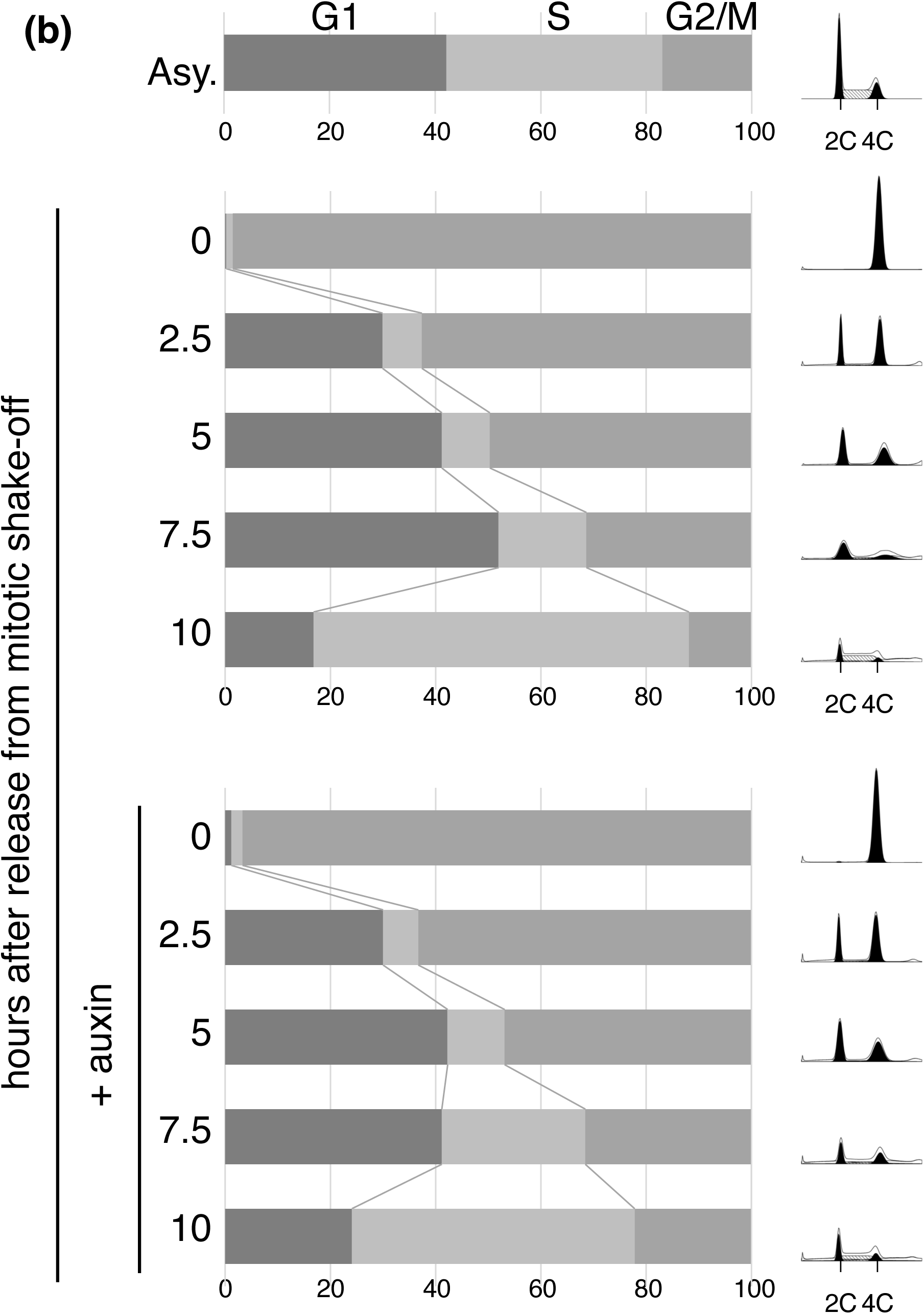
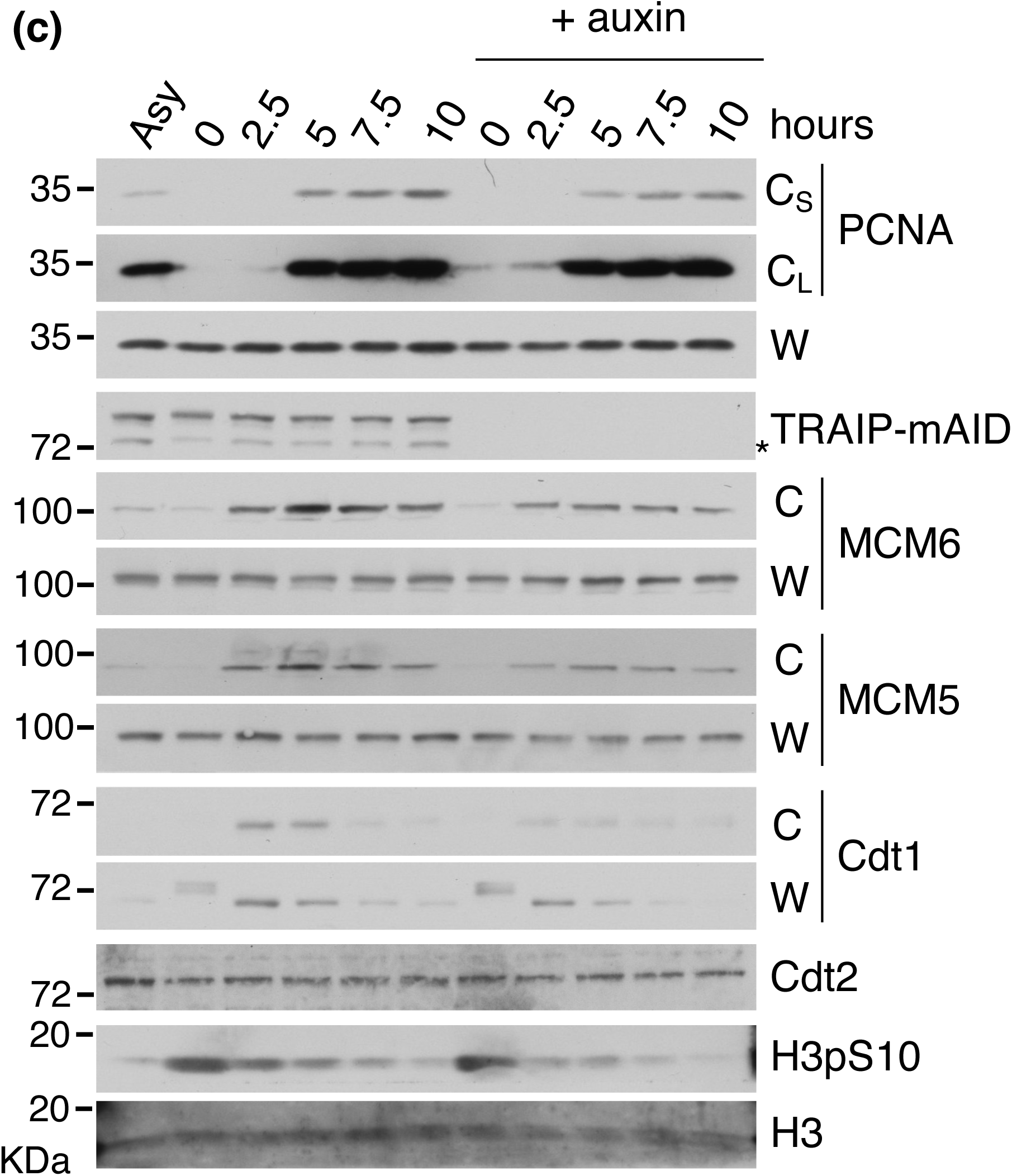
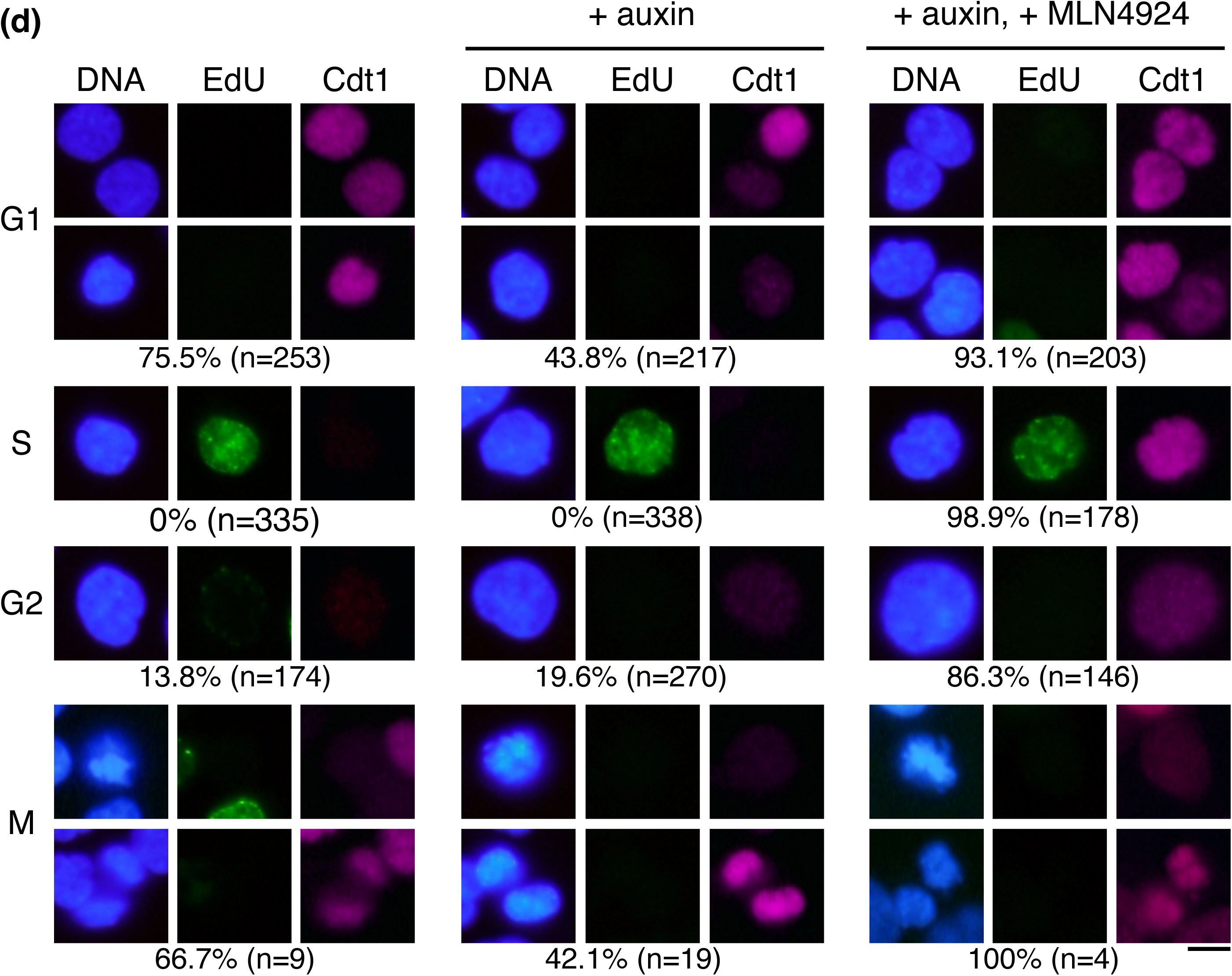
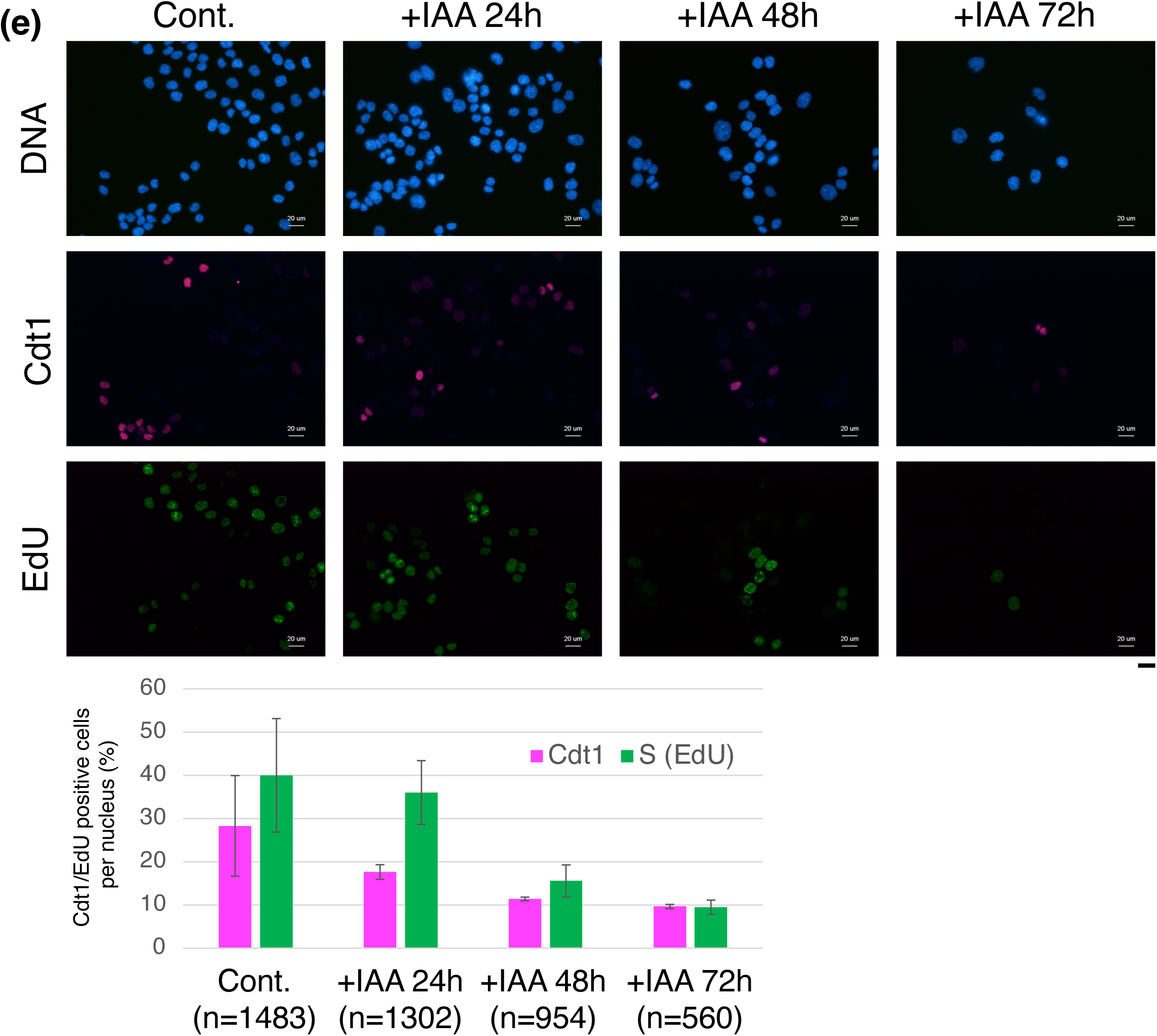
TRAIP is required for faithful DNA replication initiation. (a) TRAIP-mAID expressing cells were collected at the indicated times after the addition of auxin (5-Ph-IAA). The graph shows the ratios of cell cycle phases in asynchronously growing cells following the addition of auxin, based on the cell cycle profiles shown on the right. (b) To synchronize cells in mitosis, TRAIP-mAID expressing cells was treated by a 24-hour auxin treatment or not, followed by nocodazole for 14-hour, also in the presence or absence of auxin. Subsequently, mitotic shake-off was performed before releasing the cells, also in the presence or absence of auxin. The graphs represent the average ratios of cell cycle phases from cells arrested in mitosis and subsequently released in the absence or presence of auxin, based on the cell cycle profiles shown on the right. The data are derived from four independent experiments, with the mean values of each phase ratio displayed. (c) The whole cell extract (W) and chromatin fraction (C) of the same cell samples collected in (b) were immunoblotted with the indicated antibodies. C_S_ and C_L_ represent short exposure and long exposure, respectively, of the chromatin fraction. The band indicated by an asterisk (*) is absent following auxin treatment and, based on its molecular mass, likely represents a truncated form of TRAIP-mAID-Clover lacking the Clover tag. (d) Asynchronous TRAIP-mAID expressing cells grown in the absence or presence of auxin for 24 hours were immuno-stained with anti-Cdt1 antibody (cyan). To test the expression levels of Cdt1 during G1 phase, we excluded S phase cells based on 15 minute EdU incorporation (green) and mitotic cells based on their chromosomal structure (blue). Additionally, among EdU unincorporated cells, we also excluded G2 phase cells based on their large nucleus size. To inhibited the CRL4-Cdt2 activity, after treating the cells with auxin for 24 hours, the cells were cultured for an additional 1 hour with the addition of MLN4924. The bar indicates 10 μm. (e) Asynchronous TRAIP-mAID expressing cells grown in the absence or presence of auxin for the indicated times, followed by a 15 minute EdU incorporation, and then immuno-stained for Cdt1. The graphs depict the averages and standard deviations from three independent experiments, showing the percentage of Cdt1-positive or EdU-positive cells relative to the total number of counted nuclei (n). A decrease in both Cdt1- and EdU-positive cells, as well as a reduction in cell proliferation, was observed with increasing auxin treatment time. The bar indicates 20 μm.

To investigate whether TRAIP depletion inhibits S phase entry, cells were arrested in mitosis by treating with or without auxin for 24 hours to deplete TRAIP-mAID, followed by a 14-hour nocodazole treatment to avoid potential irreversible effects. Mitotic cells were then isolated by shake-off before release into the cell cycle. Flow cytometry analysis revealed no significant differences in cell cycle distribution between TRAIP-depleted and control cells up to 7.5 hours post-release (Figure 3b and S10). However, at 10 hours post-release, a statistically significant difference was observed in the S phase population: approximately 70% of control cells had entered S phase, compared to only 50% of TRAIP-depleted cells. Analysis of protein levels in these cells revealed that PCNA was undetectable on chromatin in nocodazole-arrested mitotic cells (time 0) in the absence of auxin (Figure 3c). However, chromatin-bound PCNA was observed in the presence of auxin, consistent with the findings presented in Figure 2c. Notably, TRAIP depletion led to a reduction in protein levels of Cdt1 and chromatin-bound MCM proteins, as observed 2.5 - 7.5 hours post-release from mitosis.

These results were confirmed by immunofluorescence microscopy of Cdt1 in asynchronously growing cells (Figure 3d). In G1 phase cells, Cdt1 was detected in about 76% of cells expressing TRAIP-mAID. Conversely, in TRAIP-mAID depleted cells after auxin treatment for 24 hours, Cdt1 signals were reduced during G1 phase to about 44%. Next, we were adding the MLN4924, an inhibitor of NEDDylation that inhibits the function of CRL4-Cdt2, after auxin treatment. As evidence of inhibited CRL4-Cdt2 activity, Cdt1 is expressed even in S phase, where it should normally be degraded, and Cdt1 expression was rescued in G1 phase even in the presence of auxin. Additionally, auxin treatment for up to 72 hours in asynchronously growing cells confirmed that the reduction of Cdt1 and EdU positive cells demonstrated that TRAIP depletion leads to Cdt1 degradation and impaired S phase progression (Figure 3e). Collectively, in TRAIP-depleted cells, PCNA remains bound to chromatin beyond S phase, leading to a prolonged G2 phase. Subsequently, this persistent chromatin-bound PCNA induces a reduction in Cdt1 expression levels during G1 phase, thereby impairing pre-RCs assembly and inhibiting the initiation of DNA replication.

Cdt1 degradation occurs via CRL4-Cdt2 during S phase, requiring chromatin-bound PCNA. In G1 phase, PCNA binds chromatin for DNA repair only upon DNA damage, triggering Cdt1 degradation through the same pathway (Nishitani and Lygerou 2004). Our previous study showed that when UV-irradiated mitotic cells progressed into G1 phase, PCNA loading onto chromatin for DNA repair triggered CRL4-Cdt2-mediated Cdt1 degradation, severely inhibiting DNA replication initiation (Morino et al. 2015). Consequently, it is postulated that the absence of PCNA binding to chromatin during G1 phase is prerequisite for accurate transition to the subsequent S phase, and TRAIP likely fulfills this role in normal cell cycle progression. However, the precise role of TRAIP in PCNA recognition and ubiquitination for removal from chromatin remains unclear. Additionally, the mechanism by which PCNA accumulates on mitotic chromatin solely due to TRAIP depletion, despite the presence of ATAD5-RFC, is yet to be elucidated. Our ongoing research focuses on distinguishing the roles of ATAD5-RFC and TRAIP in PCNA removal from chromatin, investigating their activation mechanisms and PCNA recognition throughout the cell cycle. We then aim to reveal how proper PCNA removal from chromatin contributes to genome stability.

## EXPERIMENTAL PROCEDURES

### Cell culture and RNAi

HCT116 and U2OS cells were cultured in McCoy’s 5A medium (16600-082; Gibco) and DMEM (08458-16; Nakalai), respectively, supplemented with 10% FBS and 1% Penicillin-Streptomycin at 37°C in 5% CO_2_. Mitotic arrest was induced with 75 μg/ml nocodazole for indicated times. For time course assays from mitosis, cells were treating with or without auxin for 24 hours, followed by a 14-hour nocodazole treatment, also with or without auxin. Mitotic cells were then isolated by shake-off before release into the cell cycle, also with or without auxin, and collected at specified timepoints for flow cytometry and immunoblotting. In RNAi experiments, two transfections of siRNA were performed 24 hours apart, followed by 48 hours of incubation, except for live-cell imaging. The siRNAs used in this study are described in the supporting information. To analyze cell cycle profiles, flow cytometry was performed with FACSMelody and FACSChorus software (Becton Dickinson), and the cell populations with various DNA contents were analyzed with ModFit LT (Verity Software House). For the colony formation assay and cell proliferation assay, 1,000 and 3,000 cells, respectively, were seeded into 60 mm dishes and cultured for the indicated time periods. The experimental methods are also described in the figure legends and supporting information.

### Establishment of HCT116 cell lines expressing mAID-ATAD5 and TRAIP-mAID

A CRISPR-Cas9 plasmid targeting the ATAD5 start codon region (CACCGGCTGTGGTACCAGGTCACG/agg) was constructed using pX330-U6-Chimeric_BB-CBh-hSpCas9 (Addgene #42230) (Ran et al. 2013). A donor plasmid containing the Hygro-P2A-mAID tag was created following Natsume et al. (Natsume et al. 2016). HCT116 cells expressing Tet-inducible OsTIR1 were co-transfected with these CRISPR-Cas9 and donor plasmids. To generate the TRAIP-mAID cell line, a CRISPR-Cas9 plasmid targeting the TRAIP stop codon region (CTCACTGTTCTCACGACCAC/agg) was constructed using pX330, along with a donor plasmid containing the mAID-Clover-Hygro tag. These were transfected into HCT116 cells expressing OsTIR1 (F74G) (Saito and Kanemaki, 2021; Yesbolatova et al., 2020, 2019). Bi-allelic insertion in isolated clones was confirmed by genomic PCR.

For protein complementation assays, TRAIP-mAID expressed cells were co-transfected with pCMV-hyPBase and PiggyBac transposon vectors containing TRAIP cDNA (WT or ΔPIP). TRAIP variants were amplified by RT-PCR using primers described in the supporting information.

### Immunoblotting, Fluorescent microscopy

For immunoblotting, whole cell extract and chromatin fraction were carried out as described previously (Shiomi and Nishitani 2013). The antibodies used in this study are described in the supporting information. Protein levels were calculated by examining band densities with ImageJ software. For immuno-fluorescent microscopy, the cells were treated as described previously (Shiomi and Nishitani 2013). To observed mitotic cells, the cells were simultaneous fixation and permeabilization using PTEMF buffer (Bhowmick et al., 2016). For live-cell imaging, the GFP-PCNA-expressing cDNA construct was kindly provided by Dr. Leonhardt (Leonhardt et al., 2000) and transfected into U2OS cells as the pCMV_NLS_EGFP-L2-PCNA plasmid and GFP-PCNA-expressing cells were selected using neomycin. The cells were then transfected with siRNA in a 35 mm glass-bottom dish (D11130H, Matsunami). After 24 hours, the medium was replaced with phenol red-free DMEM (21063-029; Gibco). Time-lapse imaging was performed every 15 minutes for 48 hours using a Leica TCS SP8 confocal microscope with an HC PL APO CS2 40x/1.30 OIL objective, in a humidified chamber at 37°C with 5% CO_2_.

### Statistical analysis

Statistical analyses for immunoblotting (Figure 1b and 1c), live cell imaging (Figure 2b) and flow cytometry (Figure S9) were performed by unpaired two-sided Student’s t-test, and the corresponding p-values are indicated in the figures. All measurements were taken from distinct samples and no repeated measurements were taken.

## Acknowledgments

We thank Y. Inoue and A. Kusaka (Univ. of Hyogo) for initial experiment of this work. We also thank T. Natsume (NIG), Y. Shimizu, K. Matsubara and M. Takahara (Univ. of Hyogo) for experimental support. This work was supported by JSPS KAKENHI (JP16K07257) to YaS, JSPS KAKENHI (JP21H0419 and JP22H04703) and JST CREST (JPMJCR21E6) to MTK, JSPS KAKENHI (JP20K06547) to HN.

## Author Contributions

YaS performed all experiments described in this report, except for the live cell imaging mechanical setup, which was done by AH. YuS and MTK performed AID strain construction. YaS conceived the study, designed the experiments, and wrote the manuscript. HN supervised the work.

