## Supporting Information for "The depletion of TRAIP results in the retention of PCNA on chromatin during mitosis, leads to inhibiting DNA replication initiation"

### Figure S1

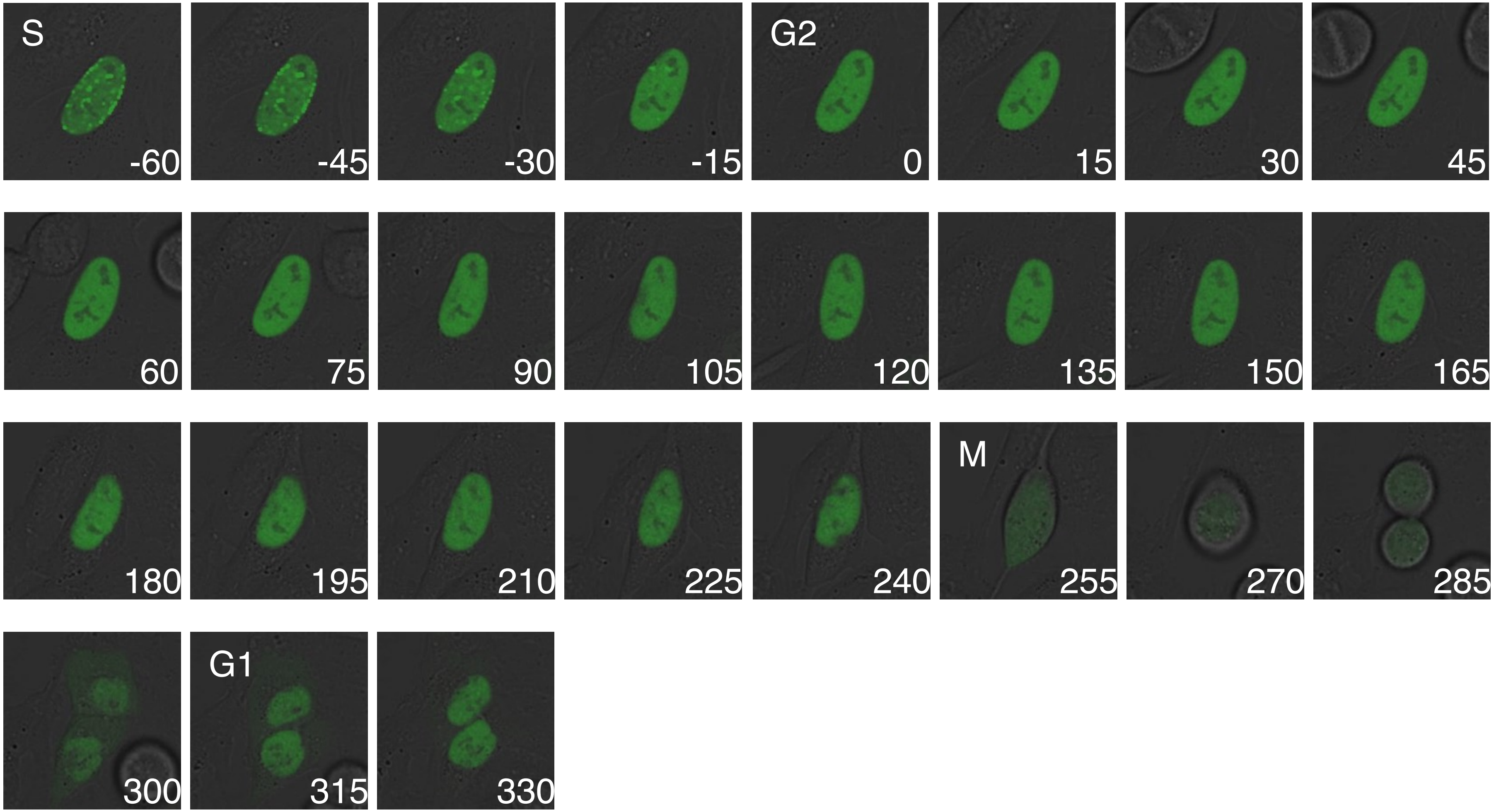


**Figure S1.** This figure presents representative time-lapse imaging of control cells. The time point at which EGFP-PCNA foci disappeared was designated as time 0. Time-lapse images are displayed at the indicated time points (in minutes). The G2 phase was defined as the interval between the disappearance of PCNA foci and nuclear envelope breakdown (NEB), while mitosis was characterized as the period from NEB to cytokinesis completion.

**Figure S2**

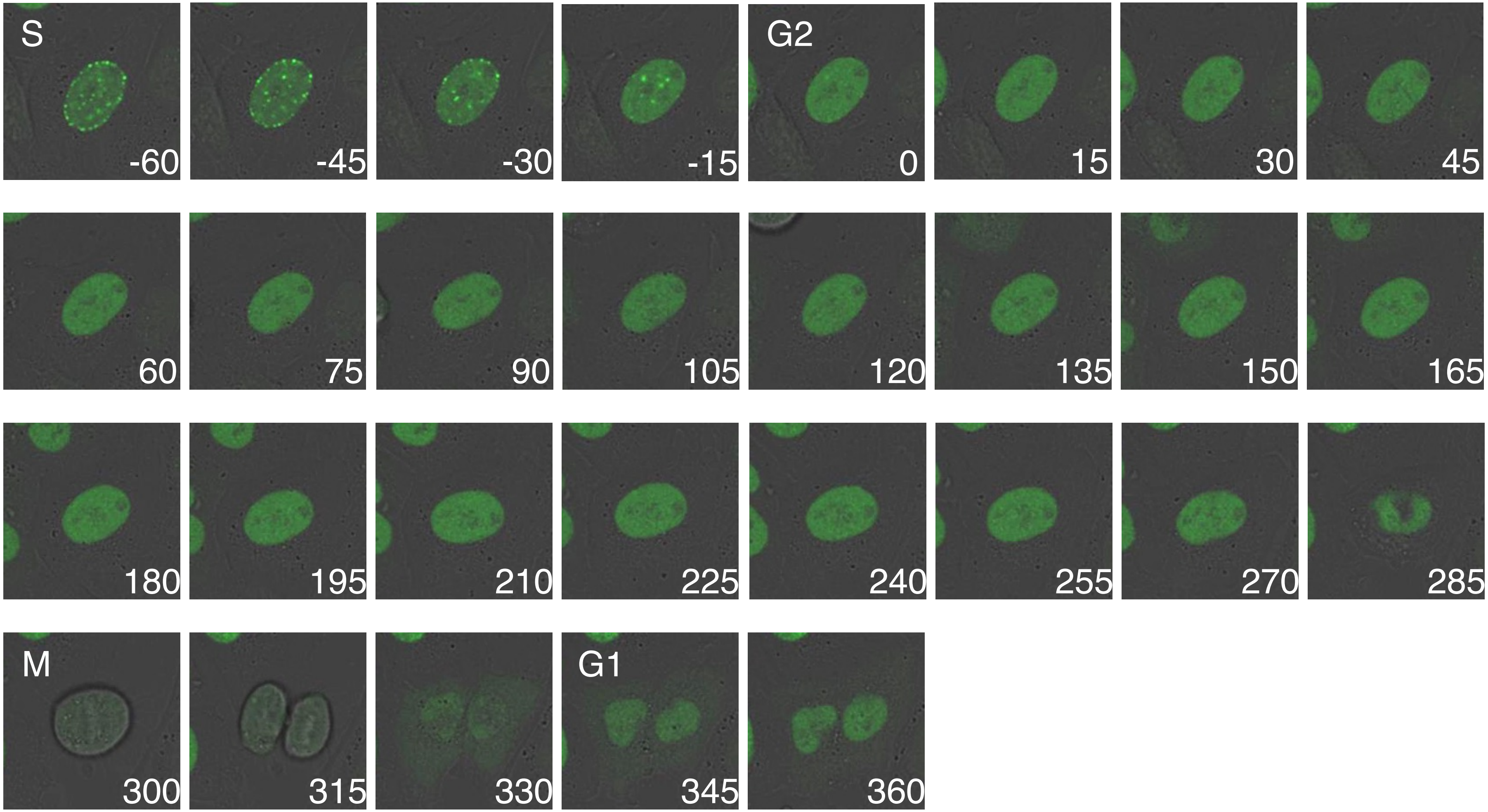


**Figure S2.** This figure presents representative time-lapse imaging of TRAIP-depleted cells by RNAi. The time point at which EGFP-PCNA foci disappeared was designated as time 0. Time-lapse images are displayed at the indicated time points (in minutes). The G2 phase was defined as the interval between the disappearance of PCNA foci and nuclear envelope breakdown (NEB), while mitosis was characterized as the period from NEB to cytokinesis completion.

### Figure S3

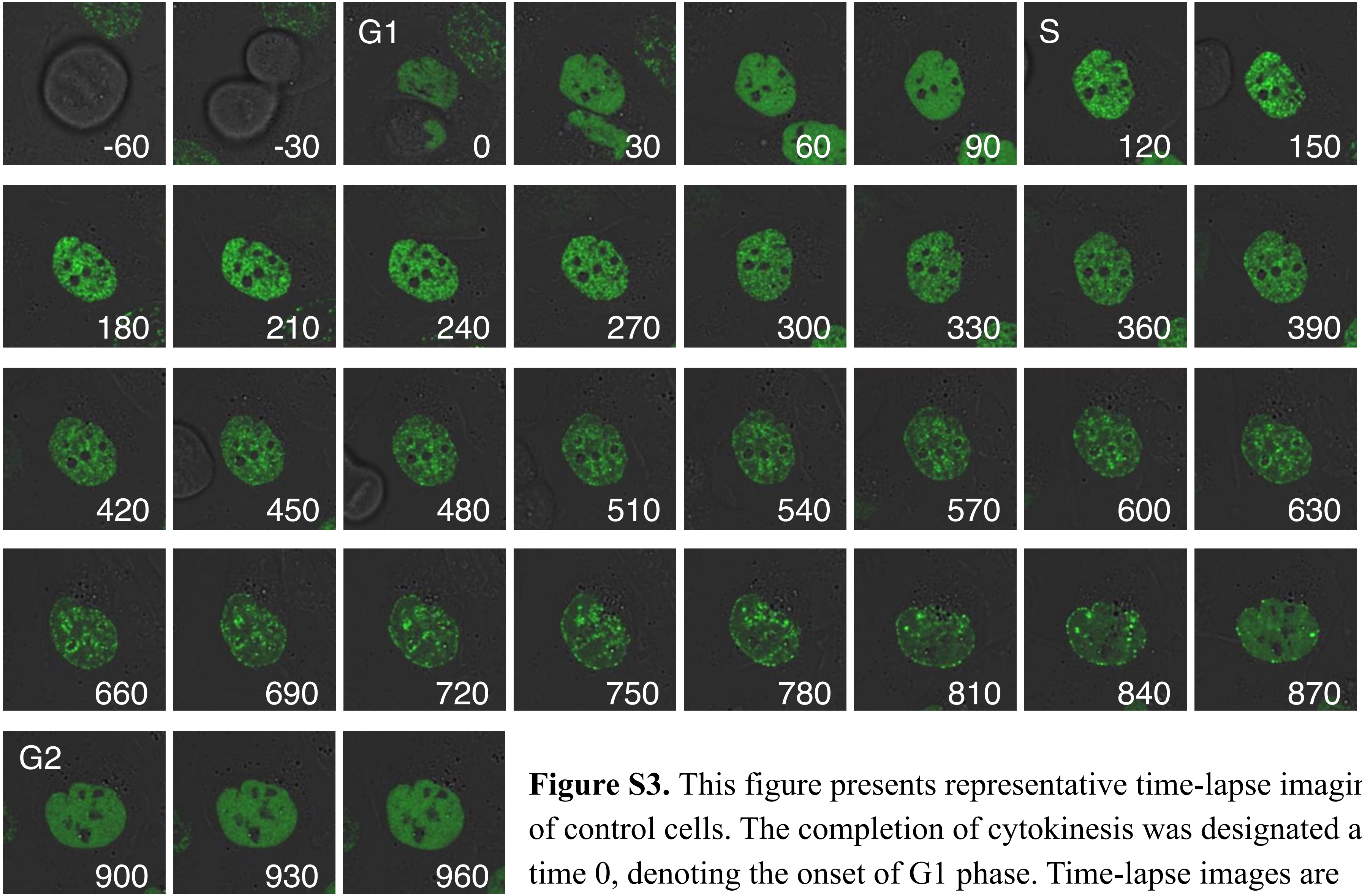


**Figure S3.** This figure presents representative time-lapse imaging of control cells. The completion of cytokinesis was designated as time 0, denoting the onset of G1 phase. Time-lapse images are presented at the indicated time points (in minutes). S phase was defined by the emergence of PCNA foci, while G2 phase was characterized by their subsequent dissolution.

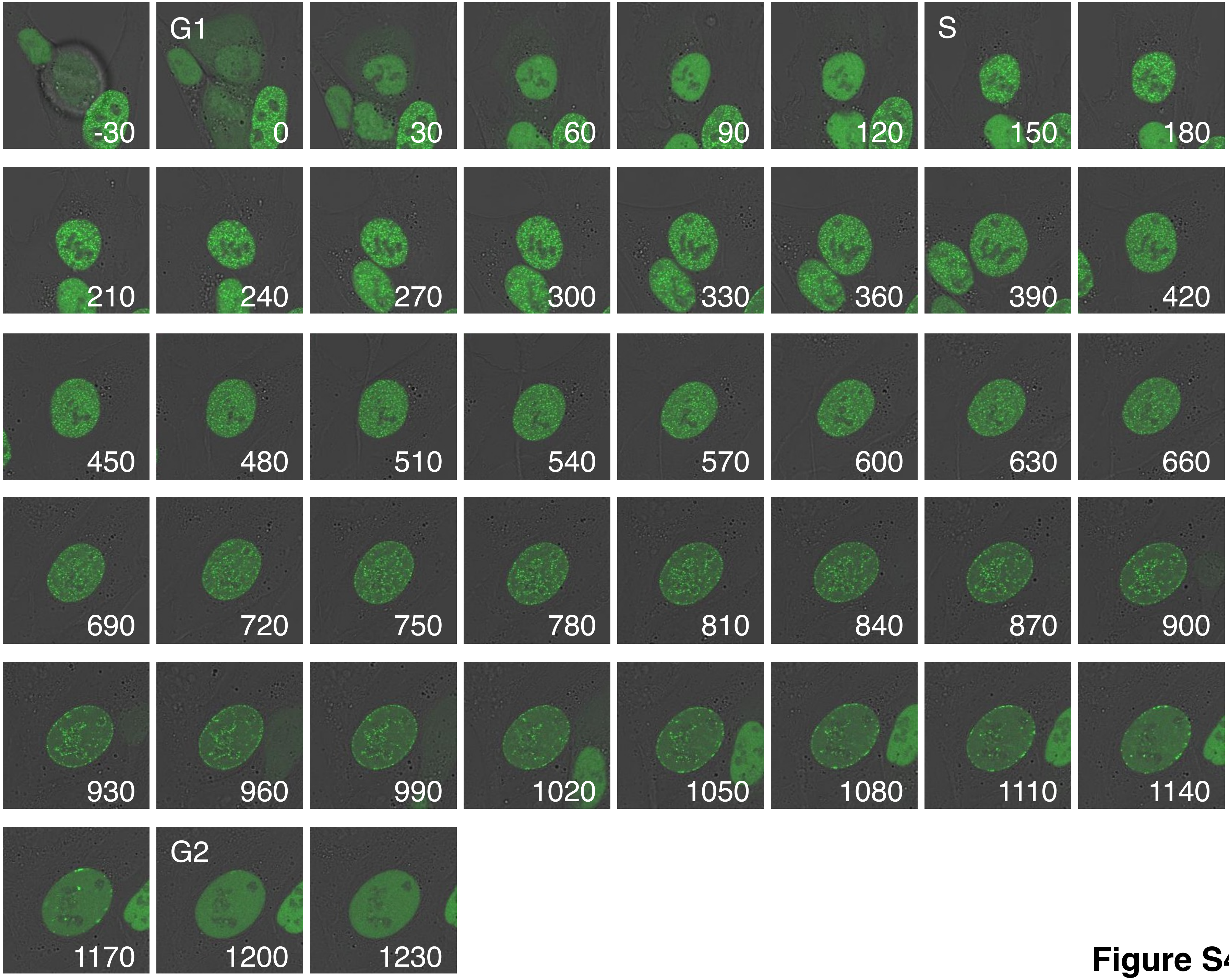
G1
S
G2
-30
0
30
60
90
120
150
180
210
240
270
300
330
360
390
420
450
480
510
540
570
600
630
660
690
720
750
780
810
840
870
900
930
960
990
1020
1050
1080
1110
1140
1170
1200
1230

**Figure S4**

**Figure S4.** This figure presents representative time-lapse imaging of TRAIP-depleted cells by RNAi. The completion of cytokinesis was designated as time 0, denoting the onset of G1 phase. Time-lapse images are presented at the indicated time points (in minutes). S phase was defined by the emergence of PCNA foci, while G2 phase was characterized by their subsequent dissolution.

#### Figure S5

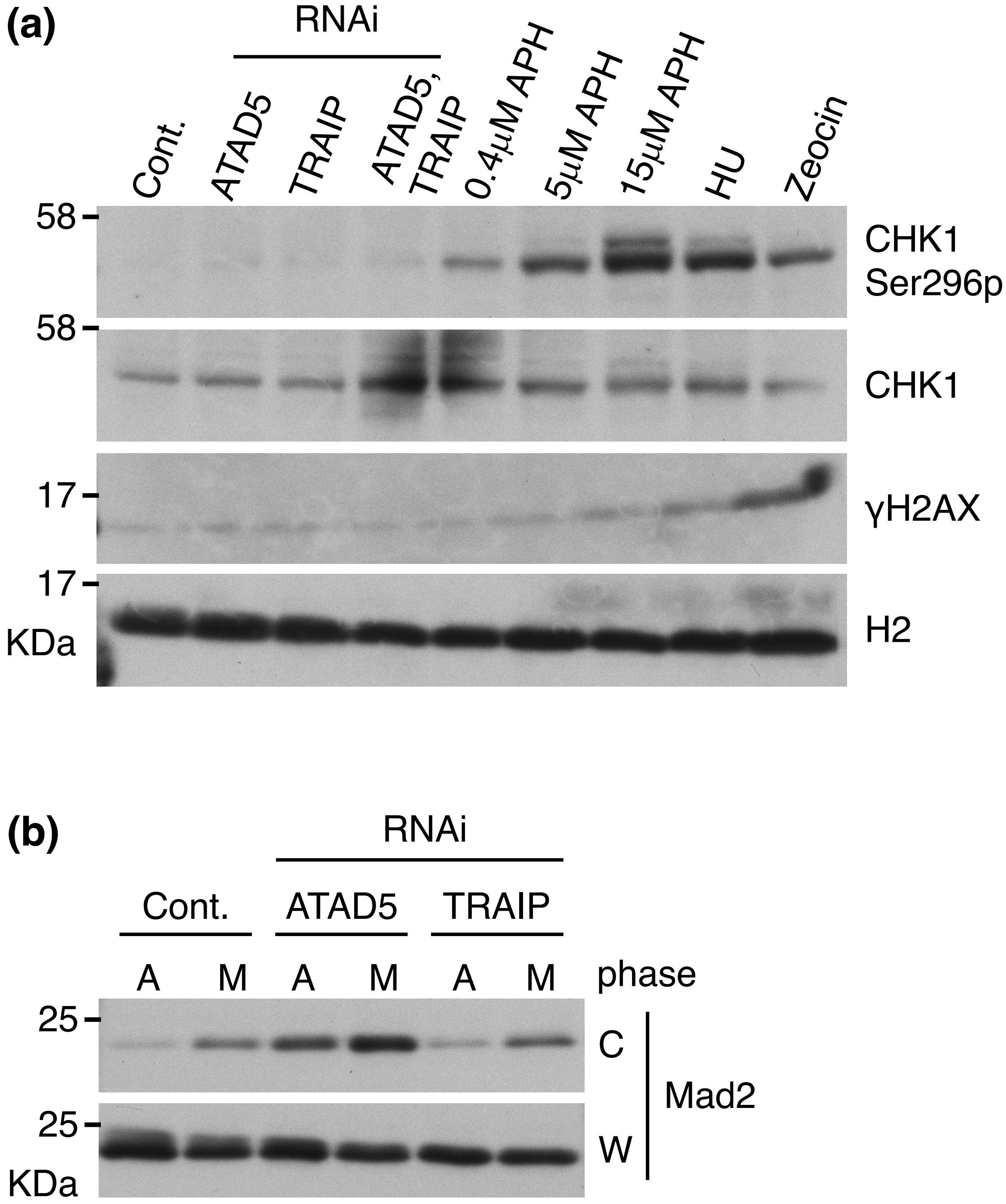
(a)
RNAi
Cont.
ATAD5
TRAIP
ATAD5, TRAIP
0.4μM APH
5μM APH
15μM APH
HU
Zeocin
58
58
17
17
KDa
CHK1 Ser296p
CHK1
γH2AX
H2
(b)
RNAi
Cont.
ATAD5
TRAIP
A
M
A
M
A
M
phase
25
25
KDa
C
W
Mad2

**Figure S5. (a)** U2OS cells were grown asynchronously and depleted of the indicated proteins by RNAi or treated with the indicated reagents (APH: aphidicolin, 1 mM HU: hydroxyurea, and 2 μg/ml Zeocin). The cells were then collected and immunoblotted with the indicated antibodies. **(b)** Asynchronously growing (A) and mitosis-arrested (M) U2OS cells, depleted of the indicated proteins by RNAi, were collected and divided into whole cell extract (W) and chromatin fraction (C), then immunoblotted with the indicated antibodies.

### Figure S6

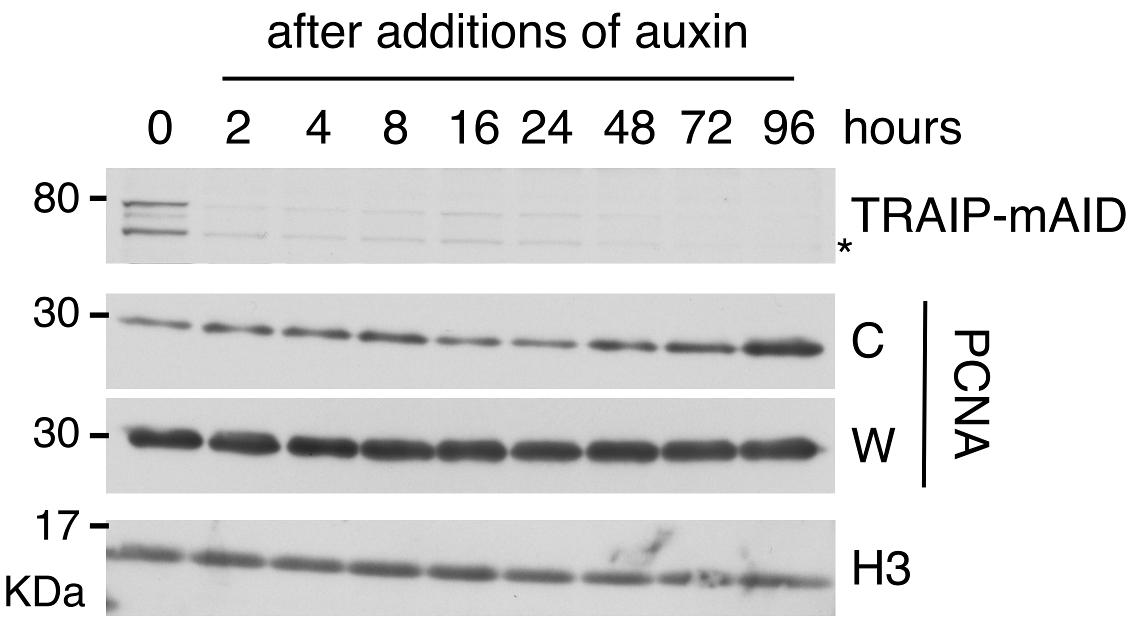


**Figure S6.** TRAIP-mAID-Clover expressing cells were collected at the indicated times after the addition of 1 mM auxin (5-Ph-IAA). The whole cell extract (W) and chromatin fraction (C) were prepared and immunoblotted with the indicated antibodies. TRAIP-mAID was almost completely degraded at 2 hours after auxin (5-Ph-IAA) addition. The band indicated by an asterisk (*) is absent following auxin treatment and, based on its molecular mass, likely represents a truncated form of TRAIP-mAID-Clover lacking the Clover tag.

Figure S7

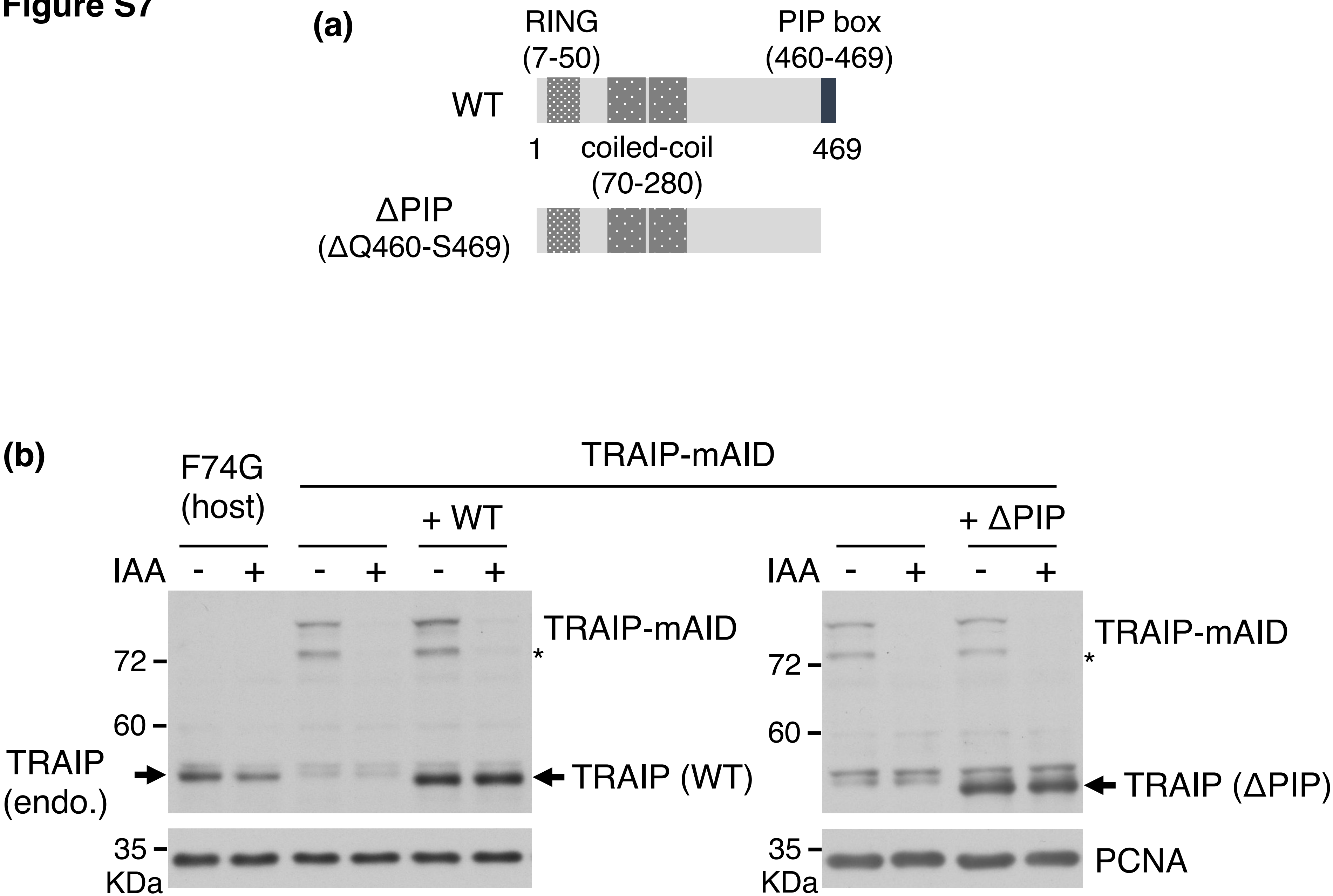
(a)
RING
(7-50)
PIP box
(460-469)
WT
1
coiled-coil
(70-280)
469
ΔPIP
(ΔQ460-S469)
(b)
F74G
(host)
TRAIP-mAID
+ WT
+ ΔPIP
IAA
- + - + - +
IAA
- + - +
72
60
35
KDa
TRAIP-mAID
*
TRAIP (WT)
TRAIP (ΔPIP)
TRAIP
(endo.)
PCNA

**Figure S7. (a)** Schematic structure of human TRAIP. The mutated site of TRAIP (ΔPIP) is indicated on the left. **(b)** Whole cell extracts of asynchronously growing F74G host cells, TRAIP-mAID expressing cells, and TRAIP-mAID expressing cells with exogenously expressed TRAIP (WT and ΔPIP) were immunoblotted with anti-TRAIP antibody. TRAIP-mAID levels decreased upon the addition of auxin (IAA). The band indicated by an asterisk (*) is absent following auxin treatment and, based on its molecular mass, likely represents a truncated form of TRAIP-mAID-Clover lacking the Clover tag.

### Figure S8

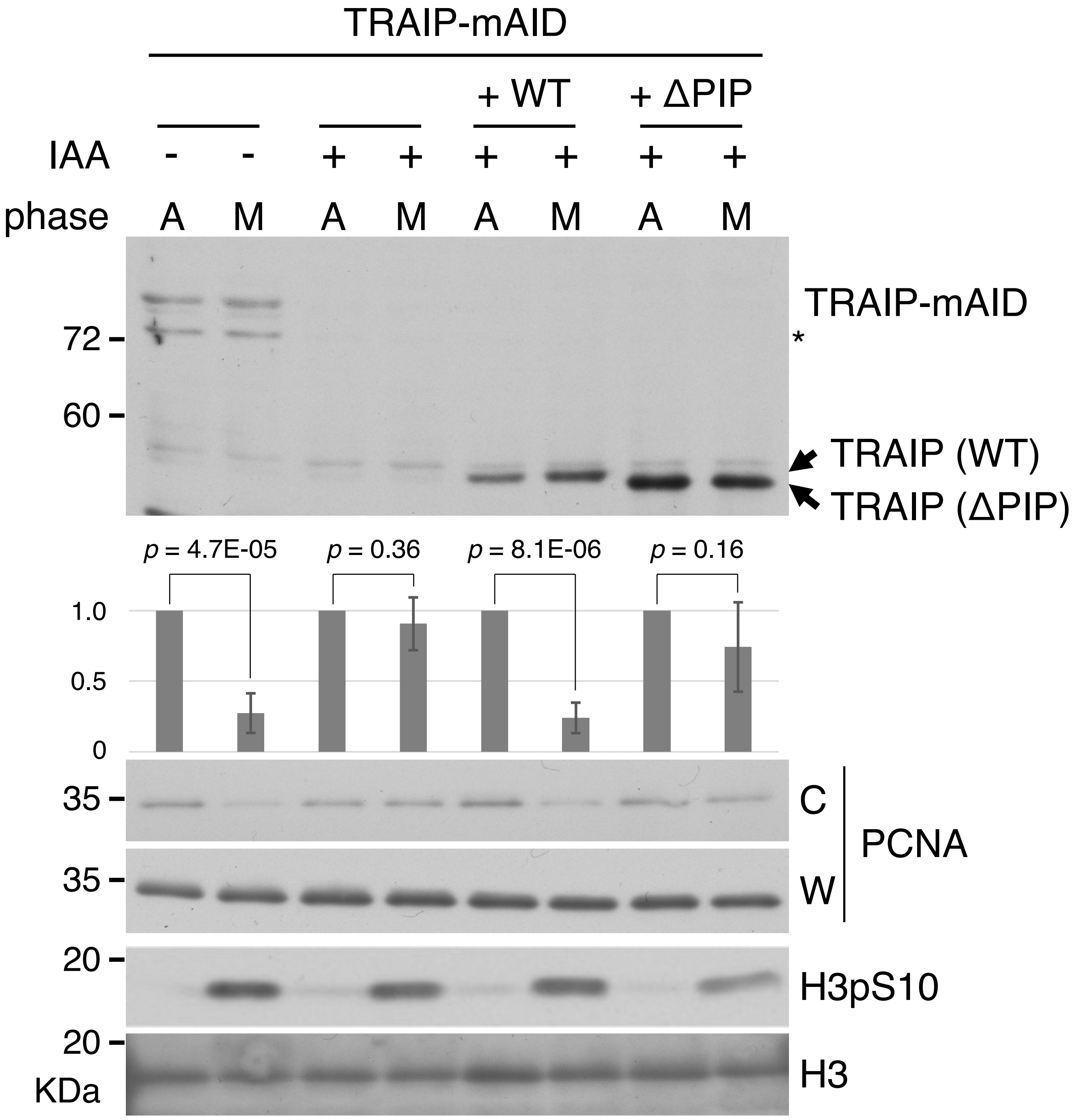

TRAIP-mAID
+ WT
+ ΔPIP
IAA - - + + + + + +
phase A M A M A M A M
TRAIP-mAID
72
*
60
TRAIP (WT)
TRAIP (ΔPIP)
p = 4.7E-05
p = 0.36
p = 8.1E-06
p = 0.16
1.0
0.5
0
35
C
PCNA
35
W
20
H3pS10
20
H3
KDa

**Figure S8.**

TRAIP-mAID expressed cells with and without introduced untagged TRAIP cDNA constructs (wild-type and PIP box deletion mutant; Figure S7). These cells were subjected to a 24-hour auxin treatment or left untreated, followed by a 24-hour nocodazole treatment (mitosis-arrested : M) or left untreated (Asynchronous : A) , in the presence or absence of auxin, were lysed and separated into W and C fractions, and the fractions were immunoblotted with the indicated antibodies. The band indicated by an asterisk (*) is absent following auxin treatment and, based on its molecular mass, likely represents a truncated form of TRAIP-mAID-Clover lacking the Clover tag. The graphs represent the averages of three independent experiments with standard deviations. The levels of chromatin-bound PCNA, normalized to total PCNA levels in the whole-cell extract (W), are displayed as vertical bars with error bars, relative to those in asynchronous control cells, which are set to 1.0. Unpaired two-sided Student's t-tests were performed, and the corresponding p-values are indicated in the figures.

#### Figure S9

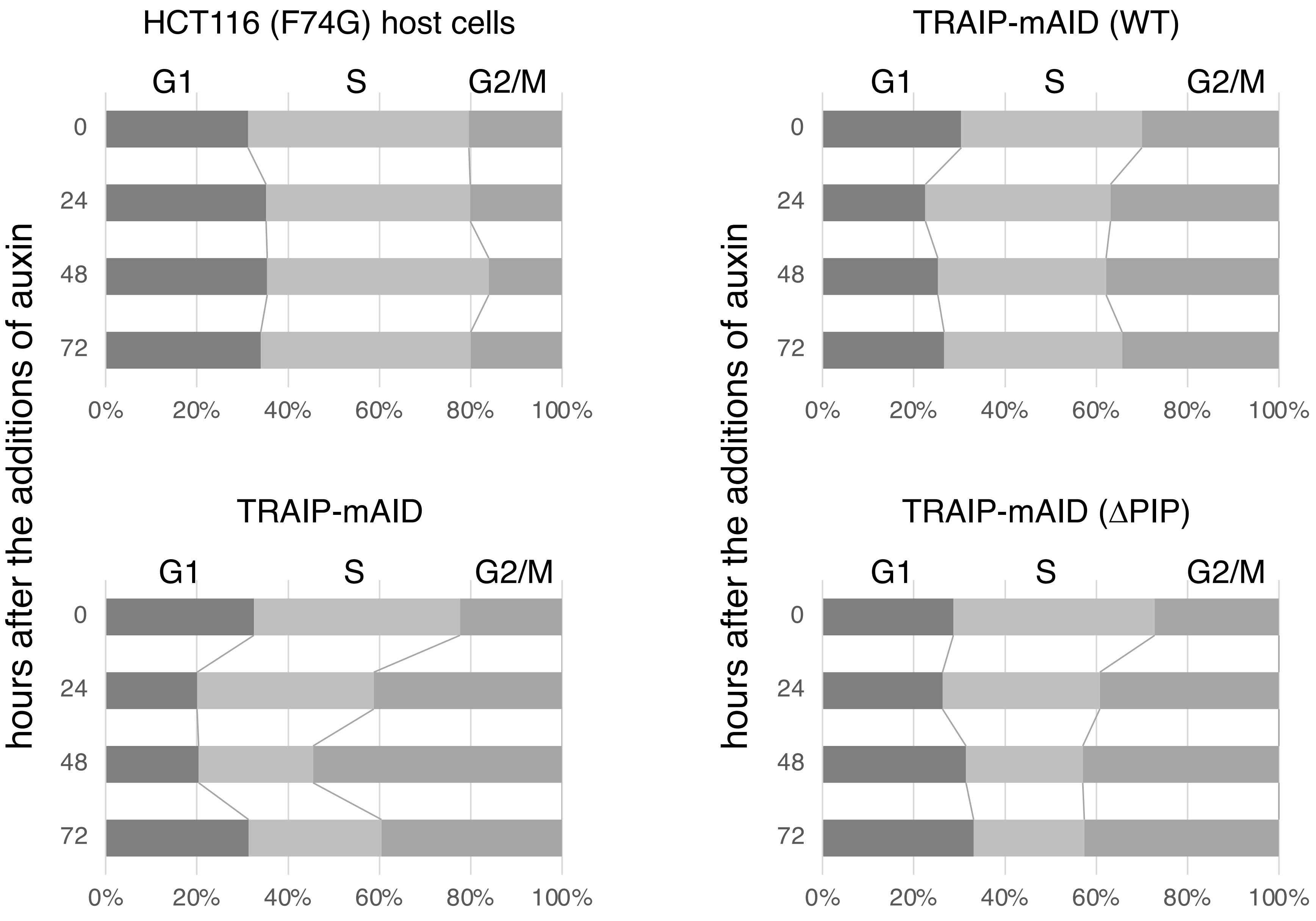
HCT116 (F74G) host cells
TRAIP-mAID (WT)
TRAIP-mAID
TRAIP-mAID (ΔPIP)
G1
S
G2/M
hours after the additions of auxin
0
24
48
72
0%
20%
40%
60%
80%
100%

**Figure S9.**

HCT116 (F74G) host cells, TRAIP-mAID expressing cells, and TRAIP-mAID expressing cells with exogenously expressed TRAIP (WT and ΔPIP) were collected 24 hours after the additions of auxin. The graphs shows the ratios of cell cycle phases in asynchronously growing cells following the addition of auxin. The data are derived from three independent experiments, with the mean values of each phase ratio displayed. Exogenously expressed TRAIP (WT) is thought to complement the function of TRAIP-mAID after auxin addition, suppressing changes in S phase populations compared to control cells. In contrast, cells expressing TRAIP (ΔPIP) exhibited cell cycle changes similar to those observed with TRAIP depletion.

### Figure S10

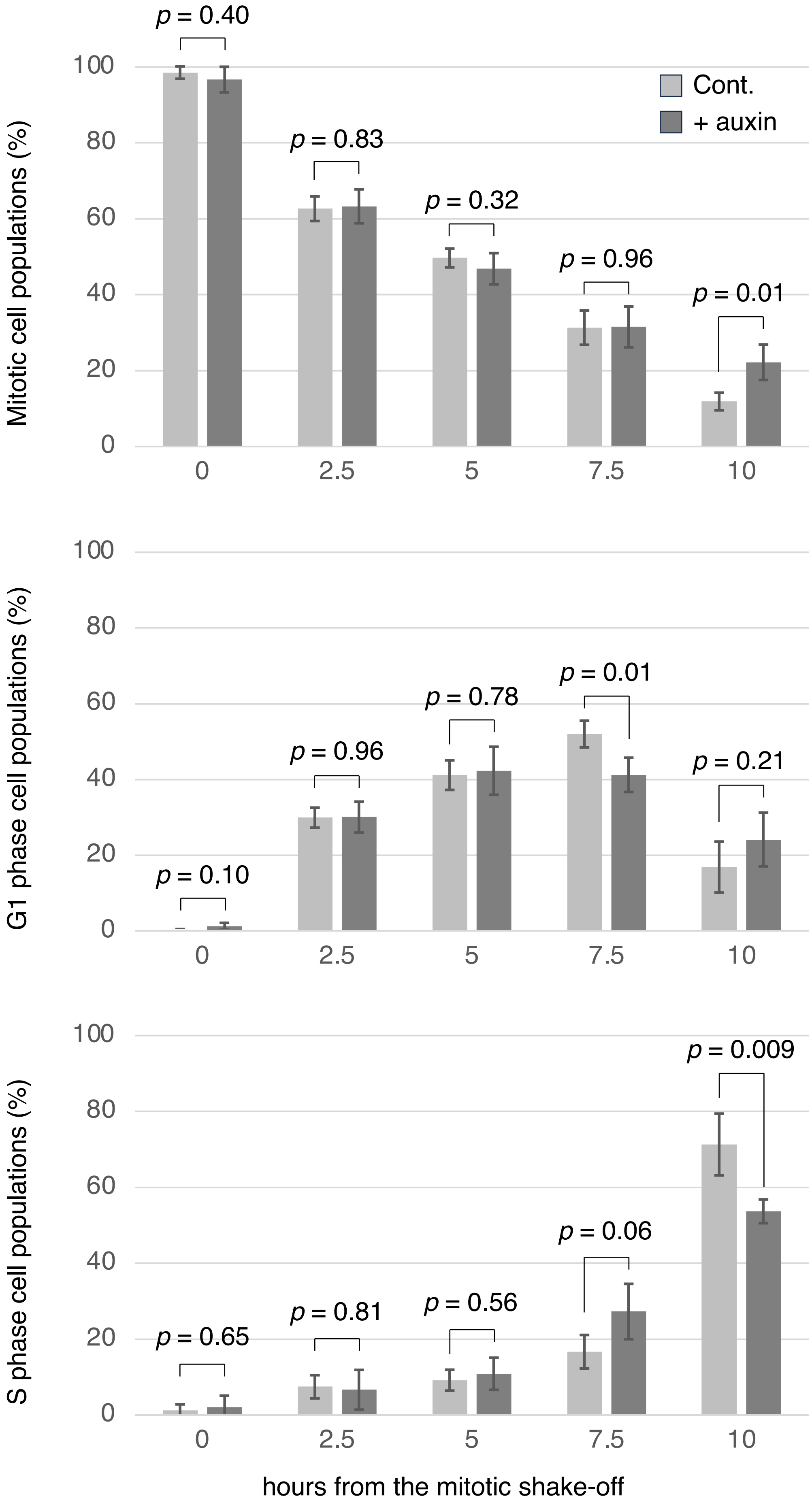

Cont.
+ auxin
Mitotic cell populations (%)
p = 0.40
p = 0.83
p = 0.32
p = 0.96
p = 0.01
100
80
60
40
20
0
0
2.5
5
7.5
10
G1 phase cell populations (%)
p = 0.10
p = 0.96
p = 0.78
p = 0.01
p = 0.21
100
80
60
40
20
0
0
2.5
5
7.5
10
S phase cell populations (%)
p = 0.65
p = 0.81
p = 0.56
p = 0.06
p = 0.009
100
80
60
40
20
0
0
2.5
5
7.5
10
hours from the mitotic shake-off

**Figure S10.** The graphs in Figure 3b has been converted into separate graphs for each cell cycle phase. The graph represents the average of four independent results, with the standard deviation. Unpaired two-sided t-tests were performed, and p-values are indicated in the figure.

#### EXPERIMENTAL PROCEDURES

##### Reagents

The final concentrations of the reagents added to the cells are as follows: 10 μM T2AA (21921; Cayman chemical), 2 μg/ml doxycycline (045-31123; Wako), 500 μM IAA (090-07123; Wako) and 1 μM 5-Ph-IAA (30-003; BioAcademia).

##### siRNA

The following siRNA were transfected at 1 μM using HiPerFect (301704; Qiagen): ATAD5 (HSS129125), TRAIP (HSS115737) from Invitrogen, and control siRNA was described previously (Shiomi and Nishitani 2013).

##### Antibodies

The following primary antibodies were used: TRAIP (AG0491), H3 (AG10644), MCM5 (11703-1-AP), MCM6 (13347-2-AP) from Proteintech, PCNA (sc-56; Santa Cruz), phospho-histone H3 (Ser10) (06-570; Sigma-Aldrich). Chk1 (2345), phospho-Chk1 (Ser 296) (2349), H2A (3636) from Cell Signaling, gH2AX (05-636; Millipore), and Mad2 (PBR452C; Covance). ATAD5 antibodies were raised in rabbits against a synthetic peptide containing aa 183 to aa 210 of human ATAD5. Antibodies for Cdt1, Cdt2 and MCM2 have been described previously (Nishitani et al. 2001; Nishitani et al. 2014). The following secondary antibodies were used: anti-mouse IgG (NA0310V) and anti-rabbit IgG (NA9340V) from GE Healthcare for immunoblotting, and: anti-mouse antibody (A11032) or anti-rabbit antibody (A11037) from Molecular Probes for immuno-fluorescence.

##### Fluorescent microscopy

To visualize DNA, hoechst 33342 (346-07951; Dojindo) was used. To identify S phase cells, cells treated with 10mM of EdU for 15 minutes and incorporated EdU was detected according to the instructions (C10337; Invitrogen).

**Primers**

TRAIP variants were amplified by RT-PCR using a common forward primer: ATATAGGTACCATGCCTATCCGTGCTCTGTGC and variant-specific reverse primers:

WT: ATATATGGATCCTCACGACCACAGGAAGGTGTCCAGC

ΔPIP: ATATATGGATCCTCAGAAGAGAGAAGGCACTGTC
